# NesSys: a novel method for accurate nuclear segmentation in 3D

**DOI:** 10.1101/502872

**Authors:** Guillaume Blin, Daina Sadurska, Rosa Portero Migueles, Naiming Chen, Julia A. Watson, Sally Lowell

## Abstract

Methods for measuring the properties of individual cells within their native 3D environment will enable a deeper understanding of embryonic development, tissue regeneration, and tumorigenesis. However current methods for segmenting nuclei in 3D tissues are not designed for situations where nuclei are densely packed, non-spherical, heterogeneous in shape, size, or texture, all of which are true of many embryonic and adult tissue types as well as in many cases for cells differentiating in culture.

Here we overcome this bottleneck by devising a novel method based on labelling the nuclear envelope (NE) and automatically distinguishing individual nuclei using a tree structured ridge tracing method followed by shape ranking according to a trained classifier. The method is fast and makes it possible to process images that are larger than the computer’s memory. We consistently obtain accurate segmentation rates of >90% even for challenging images such as mid-gestation embryos or 3D cultures. We provide a 3D editor and inspector for the manual curation of the segmentation results as well as a program to assess the accuracy of the segmentation.

We have also generated a live reporter of the NE that can be used to track live cells in three dimensions over time. We use this to monitor the history of cell interactions and occurrences of neighbour exchange within cultures of pluripotent cells during differentiation.

We provide these tools in an open-access user-friendly format.

## Introduction

Studying the properties of individual cells in relation to their neighbours in intact tissues is the first step towards understanding the cell-cell interactions that govern the behaviour of tissues. Single cell analysis can also reveal heterogeneity in cellular properties that is masked by lower-resolution population-averaging methods. Automated computational image analysis is a particularly attractive approach because it is free from operator-bias, provides quantitative data, and reveals sub-visual information that would not otherwise be apparent (Caicedo et al., 2017; Dufour et al., 2017; Keller, 2013; Meijering et al., 2016).

In order to meet the need for automated nuclear segmentation in 3D, a wide array of methods have been developed, reviewed in (Dufour et al., 2017; Kan, 2017; Meijering, 2012; Nketia et al., 2017). It is becoming increasingly apparent, however, that there is no single ‘one size fits all’ solution to segmentation. New solutions are needed for situations where nuclei are densely packed, non-spherical, or heterogeneous in shape, size, or texture: these things apply to many embryonic and adult tissue types as well to cells differentiating in culture. There is also a need to reduce the time and computational power required for segmentation of each cell: this becomes a limiting factor when imaging whole embryos, large tissues, when analysing time-lapse data, or in any other situation where large numbers of nuclei need to be identified. Finally, adoption of published methods by the community can be limited, highlighting the importance of creating well documented and user-friendly software (Cardona and Tomancak, 2012; Carpenter et al., 2013; Meijering et al., 2016).

Here, we report a new approach to overcome these bottlenecks in quantitative image analysis of individual cells in 3D. Rather than relying on staining for nuclear content (for example DAPI or Hoescht staining), we instead detect the nuclear envelope (NE). This makes it easier to identify individual nuclei that are in close contact with each other and does not suffer from segmentation problems associated with textured nuclear staining, unusually shaped nuclei, or cell debris. Furthermore, the NE of individual nuclei are easily discernable by eye in crowded tissues, and so manual correction of any mis-segmented nuclei becomes easier than is the case for DAPI stained nuclei. We provide a user-friendly 3D-4D editing tool to rapidly correct any segmentation errors.

We test our method alongside other previously-published user-friendly methods, with focus on mid-gestation embryos, 3D cultures of pluripotent stem cells and pluripotent stem cell-derived neural rosettes. These contexts are chosen because they exemplify many of the problems in 3D segmentation that we set out to address: crowded overlapping nuclear signals, non-spherical nuclei, large numbers of nuclei, and decreases in signal intensity towards the centre of large structures. Using our editing tool, we manually annotated thousands of nuclei in these contexts to generate a set of segmented images which are made publicly available and which could be useful as a standard segmentation benchmarking dataset (Ulman et al., 2017).

We consistently obtain accurate segmentation rates of >90% in these challenging contexts. We demonstrate the utility of this tool by measuring the expression of the transcription factor Tcf15 at single cell resolution across the anterior-posterior axis of the E8.75 mouse embryo. We also developed a non-disruptive method to fluorescently label the NE in live cells and used this to follow the history of cell-cell interactions and to document neighbour-exchange as pluripotent cells differentiate in 3D culture.

This new tool, named “NesSys (Nuclear Envelope Segmentation System), adds to the expanding toolkit of segmentation approaches, each of which has its particular strengths. NesSys is particularly well suited for segmenting large numbers of nuclei arranged in complex 3D configuration without requiring extensive amounts of time, computational power, or user-input. We provide NesSys as an open-access, well documented, easy to install and user-friendly software.

## Results

### Segmentation of nuclei in groups of densely packed cells

Nuclei are commonly detected based on fluorescent markers of nuclear content, for example dapi. However, it can become difficult to distinguish individual cells based on dapi staining when nuclei are densely crowded, for example during neural differentiation of pluripotent cells (Fig. 1A left), 3D cultures (Fig. 1A right) or in densely packed tissues *in vivo* (Fig. 1B).

**Fig. 1.**
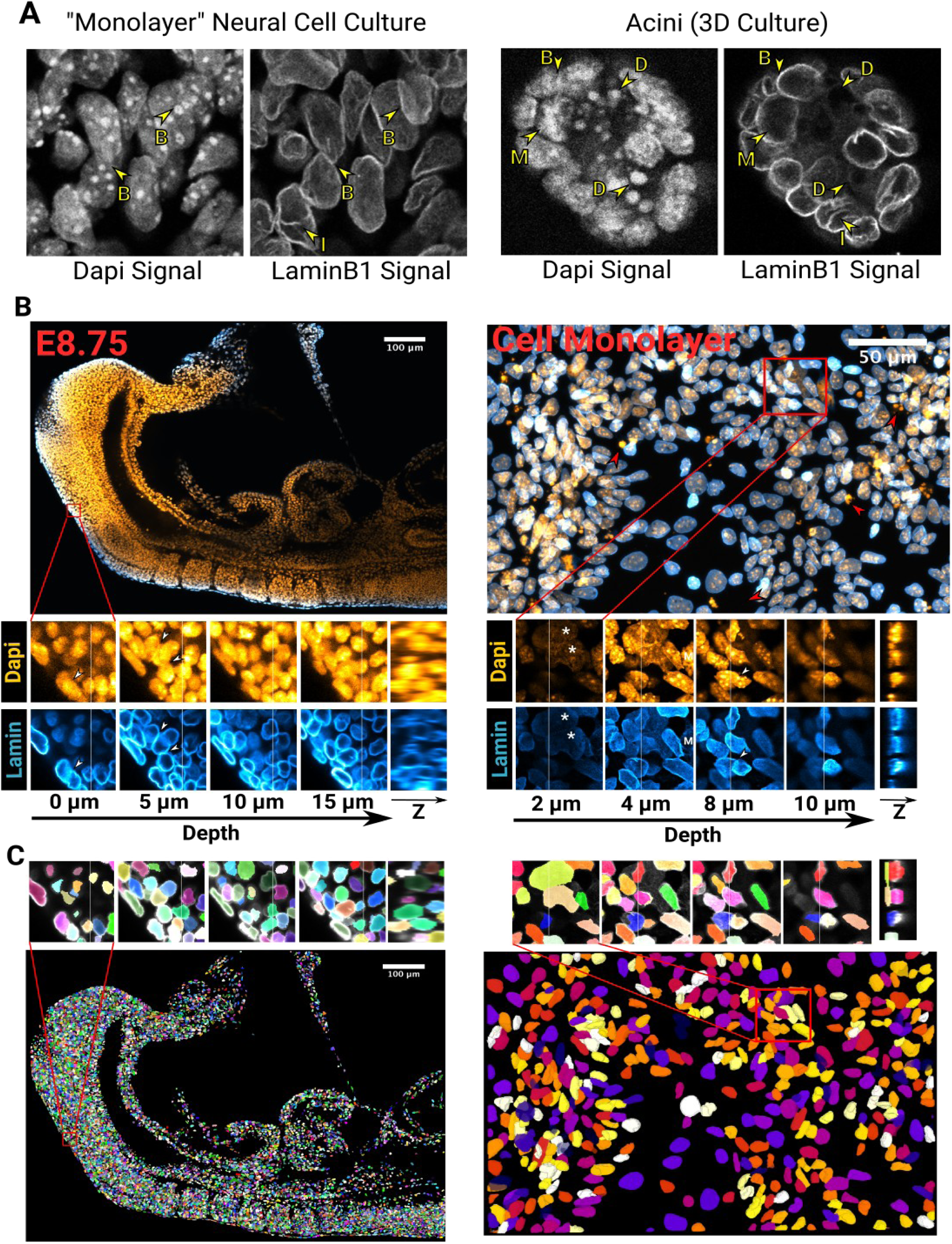
Segmentation challenges and Nessys results. **(A)** Side by side comparison of the Dapi and LaminB1 signals obtained by confocal microscopy of pluripotent stem cells differentiated into neural cells (left) or grown in a 3D matrix (right). ‘B’ arrows indicate boundaries between touching nuclei, ‘D’ arrows point at cell debris, the ‘M’ arrow shows a mitotic nucleus and ‘I’ arrows indicate invaginations of Lamin B1 into the nucleus. **(B)** Confocal micrographs showing the diversity of cell shapes and volumes found in an E8.75 mouse embryo (left) and within a ‘monolayer’ of cultured cells (right). Dapi is shown in orange and LaminB1 in cyan. For each image, a magnified region is shown as a series of planes along the z axis of the image for both the Dapi channel or the LaminB1 channel. An image constructed along the YZ axes is also shown. The faint vertical bar in the XY planes indicate the location of the YZ image. White arrows: loss of nuclei edge in the dapi signal, Red arrow: cell debris apparent in the dapi signal, White asterisks: flat and large nuclei distinct from their surrounding cells. **(C)** Images of segmented nuclei obtained with NESSys. Nuclei are assigned a unique label and a random colour. The same regions as in panel B are shown. Notice how overlaping nuclei with distinct morphology are identified accurately.

We decided to avoid this problem by using the nuclear envelope (NE) as a landmark for segmentation. Staining for Lamin B1, an intermediate filament that marks the NE (Dittmer and Misteli, 2011), allows crowded nuclei to be clearly distinguished by eye (Fig. 1). This observation prompted us to explore whether using the NE signal as an input for an automated detection method would be a successful approach.

This led us to develop a method called Nessys (Nuclear Envelope Segmentation System) that is able to reliably detect individual nuclei in crowded groups (Fig 1C: see also Movies S1, Fig. 3 and Fig. 4A, B). Below we give an overview of this method and a demonstration of its utility.

### Overview of the Nessys Segmentation Method

The main steps of the Nessys segmentation method are briefly summarised here (Fig. 2). An in depth description of complex procedures (asterisks in Fig. 2) is also available in Supplementary Materials.

- A steerable filter developed by (Jacob and Unser, 2004) is used to enhance the NE signal in each slice of the 3D image (Ridge enhancement).
- Regions of maximum fluorescence intensity are then identified using a Difference of Gaussian detector (Marr and Hildreth, 1980). These form the starting points for a dynamic ridge-tracing process, which continuously identifies and moves to the brightest adjacent pixel in an iterative process (see also Fig. S1 and Movie S2).
- Over time, this tracking algorithm identifies branch points and iteratively establishes the most likely segmentation solution using a naive Bayes classifier trained by the user (Fig. S1 and S2).
- Once this process has been completed for each z slice, 2D areas are linked together into 3D volumes. At this point, information from adjacent slices helps refining the initial 2D segmentation results (Fig. S3).

**Fig. 2.**
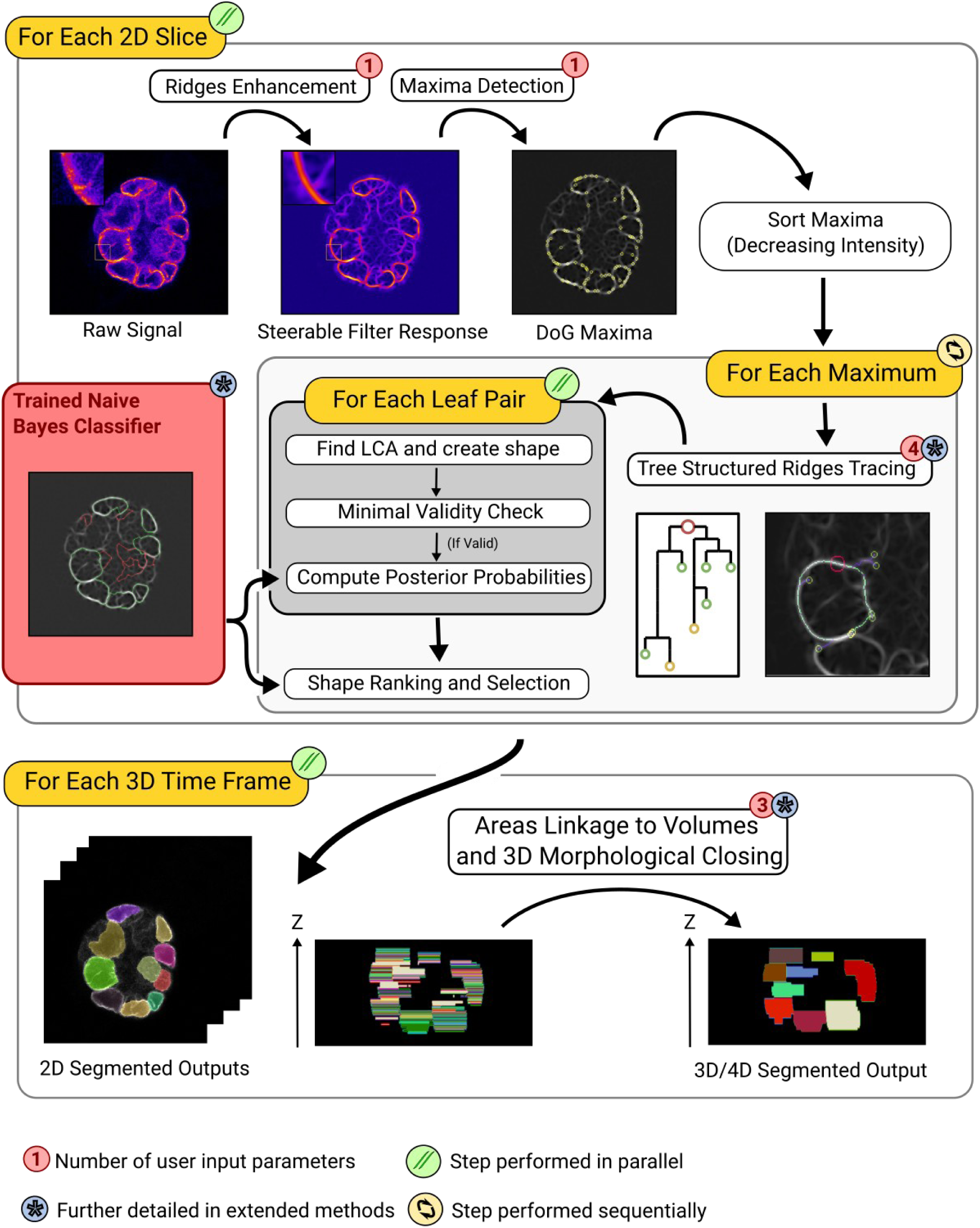
Overview of the NESSys Segmentation Method: The main steps of the method are shown in white boxes annotated with a blue asterisk whenever the step is further detailed in supplementary materials. Red numbers indicate how many user-defined parameters are required at each of these steps. Where possible, a snapshot of the intermediate output is shown. Iterations are represented with orange boxes annotated with an icon to indicate whether iterations are parallelised or sequential. Instructions which are part of the same iteration are contained within the same rounded box. In the ‘Tree Structured Ridge Tracing’ step, a full tree is overlaid on the image and corresponds to the diagram on the left of the image. The red circle represents the root (maximum where the procedure was initialised), smaller circles indicate the leaves of the tree, and lines represent the branches of the tree. The use of a (reusable) trained Naive Bayes classifier is shown with a red box. This classifier is trained by the user on a shape database before running the method. **LCA:** Least Common Ancestor, **Minimal Validity Check:** This step tests performs a minimal validity test to determine if the shape should be included in the list of shapes to be ranked by the classifier: 1) the tested shape must not overlap with an existing shape 2) simple features such as compactness or area must be within user-defined constraints.

Notably, the most computer-intensive aspects of this process are performed in parallel rather than in series: this helps to greatly reduce the computational time taken to analyse large images (see also Fig. 4D).

#### An open access utility program and dataset for the benchmarking of nuclear segmentation methods

Determining the performance of a segmentation method requires a gold standard (or Ground Truth - GT) image dataset (Amat et al., 2014; Coelho et al., 2009; Maška et al., 2014; Ulman et al., 2017). GT datasets may be computer-generated (Rajaram et al., 2012; Svoboda and Ulman, 2017; Wiesmann et al., 2017), or may consist of images which have been manually segmented by biologist specialists (Coelho et al., 2009). Recent initiatives have undertaken to make such datasets publicly available (Ulman et al., 2017). However, currently available GT datasets for nuclear segmentation only contain a ‘nuclear content’ signal and in most cases do not encompass the diversity in nuclear shape and sizes or the level of complexity found in mammalian systems.

For these reasons, we have generated a new image dataset with thousands of manually annotated nuclei for segmentation benchmarking. We used both a NE signal (LaminB1) and the more conventional chromatin signal (Dapi) (Fig. 3 and Table S1).

**Fig. 3.**
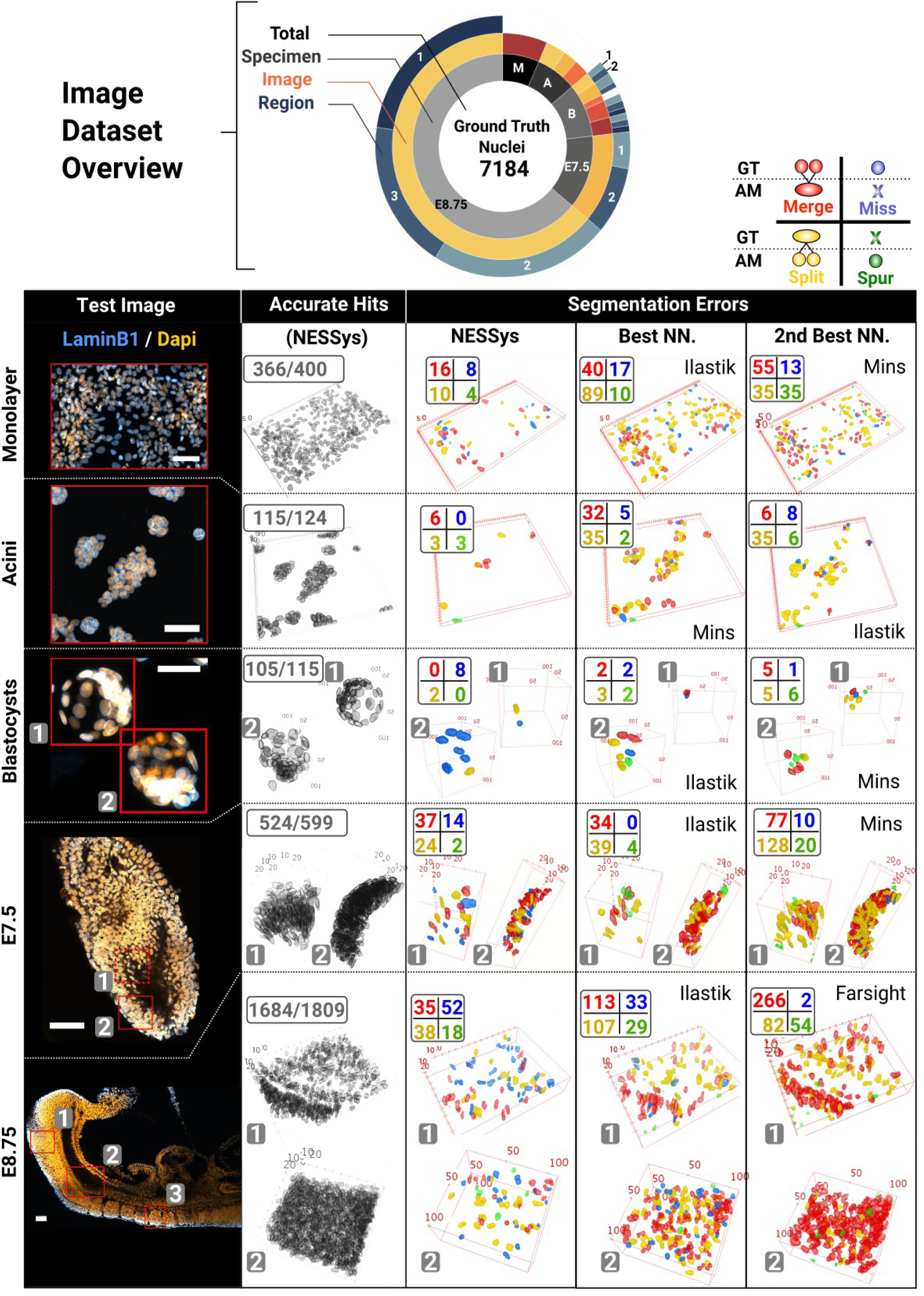
Overview of the manually segmented image dataset and sample error maps created with Nessys’ validation and benchmarking tools. The sunburst chart at the top of the figure shows the organisation of the image dataset. The size of the rings sections is proportional to the number of manually segmented nuclei (ground truth nuclei) within the dataset. (Grey divisions: biological specimen, M: Monolayer, A: Acini, B: Blastocysts. Orange divisions: images, blue divisions: manually segmented regions. NB: no sub-division is drawn when the image was manually segmented in full). Representative test images are shown in the left column of the table below, **Scale Bars:** 100 μm. For each image, red outlines represent the manually segmented image regions used to evaluate accurate hits and errors which are shown as 3D maps in the other columns. If multiple regions are drawn, a number in a grey box indicates the correspondence between a given region and the matching 3D map and sunburst section. 2^nd^ column: accurate nuclei identified with Nessys. 3^rd^ column: Maps of Nessys errors. 4^th^ and 5^th^ column: Map of errors for the best and 2^nd^ best non-Nessys (**NN**.) methods. The legend for these errors is indicated in the top right corner of the figure. GT: Ground Truth, **AM:** Automated Method, **Merge:** Only one nucleus found in AM when several nuclei are present in GT, **Miss:** nucleus found in GT but absent from AM, **Split:** several nuclei found in AM when only one is present in GT, **Spur.:** Spurious nucleus found in AM which does not exist in GT. Please note that the Monolayer and the E8.75 images are the same images as in Fig. 1B.

This dataset includes images with crowded, overlapping, heterogeneous, non-spherical nuclei in several different contexts, representing the particular segmentation challenges that we have set out to address (Fig. 3). Images include:

- High density monolayer culture of differentiating mouse ES cells
- 3D cultures of differentiating mouse ES cells embedded in matrigel
- E3.5 mouse blastocysts
- Gastrulating mouse embryo at E7.5, focusing on a region of the distal epiblast
- Post-gastrulation mouse embryo at E8.75, focusing on three regions of the embryo which capture both epithelial and mesenchymal structures as well as variations in signal intensity due to light scattering or antibody penetration issues.

We called this dataset DISCEPTS (Differentiating Stem Cells & Embryos are a Pain To Segment) and we intend to deposit this dataset on a publicly available repository which will then be linked to from our website [https://framagit.org/pickcellslab/nessys].

In order to perform both manual annotation of nuclei in 3D and segmentation benchmarking, we developed additional programs that we provide alongside the Nessys method (Fig. S4):

1. a 3D painting tool to establish GT segmentation data (we estimate that ~100 3D nuclei were manually segmented per hour). Note that this tool can also be used as a visual inspector and 3D editor to curate outputs from any automated methods.
2. An extensible segmentation comparator which can compute the benchmarking metrics described in (Coelho et al., 2009) as well as 3D maps of accurate hits and segmentation errors (Fig. 3 and S4). This makes it possible to visually compare the frequency and distribution of each class of segmentation error for each tested segmentation output.

Using these new tools and our DISCEPTS dataset, we compared the performance of Nessys to other previously published popular methods (Fig. 3 and 4 and Materials and Methods) listed below:

- Mins (Lou et al., 2014): originally designed for detecting cells within preimplantation embryos based on nuclear content.
- Farsight (Al-Kofahi et al., 2010): designed for detecting nuclei based on nuclear content.
- Ilastik (Sommer et al., 2011): A more generalist method which uses machine learning. NB: to harness the full Ilastik capability and obtain the best performance we used both the NE and the dapi channels as input to inform the Ilastik method (See also Methods).

**Fig. 4.**
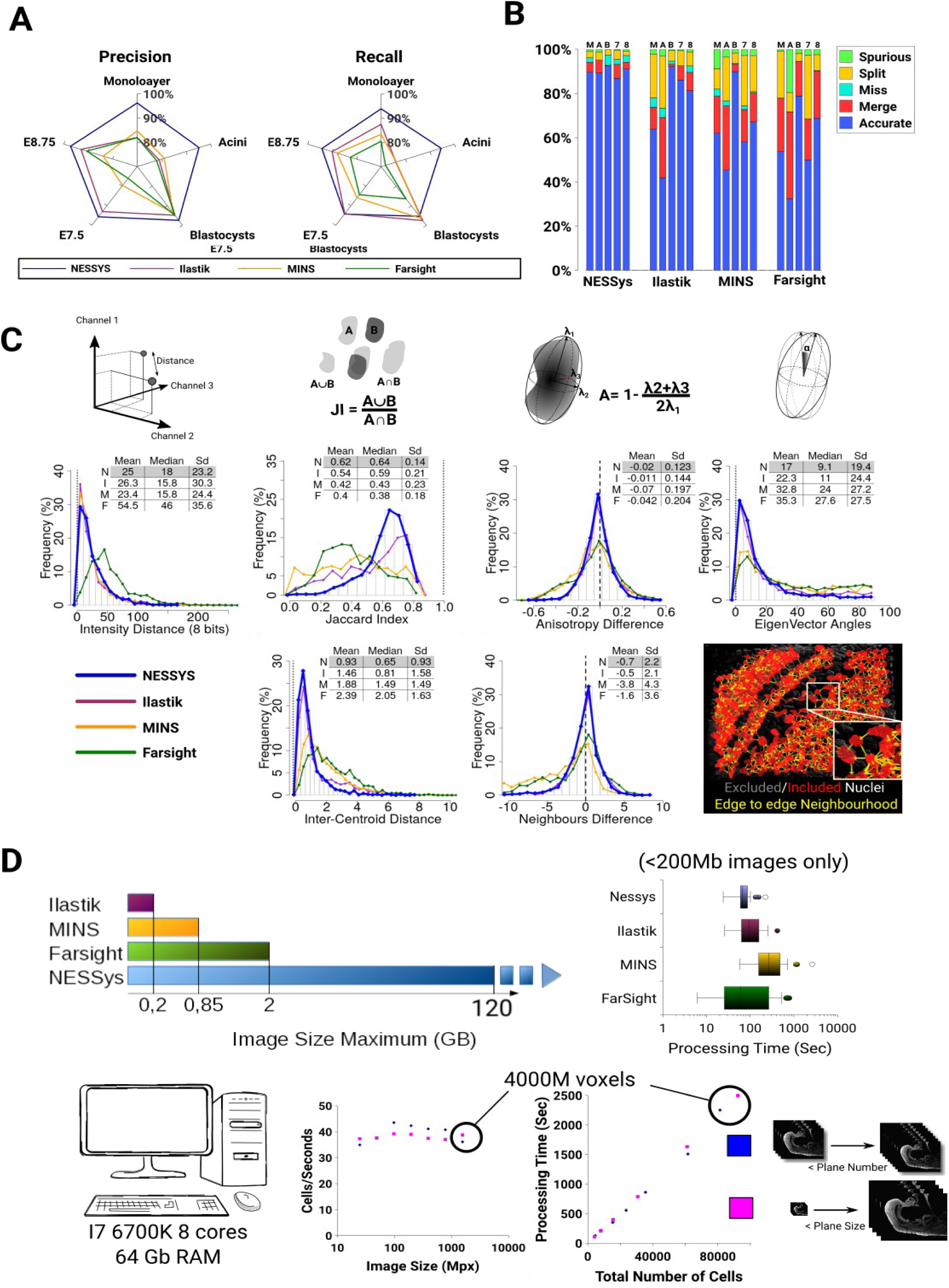
Error counting, morphological accuracy and computational time benchmarking. **A** Radar charts representing precision and recall for each tested method over the DISCEPTS dataset. **B** Stacked bar chart showing the proportion of detection accuracy and errors for each tested segmentation method (M: Monolayer, A: Acini, B: Blastocysts, 7: E7.5, 8: E8.75). **C** Histograms of the deviation of morphological features from the ground truth (to create a single graph for each feature, 200 cells from each biological dataset were randomly sampled and pooled together so each each dataset would contribute equally to the graph. A table summarises the mean and standard deviation of the distribution for each tested method (N: Nessys, I: Ilastik, M: MINS, F: Farsight). The vertical dashed line shows the value of the measure for the ground truth. An illustration is also given for most features to illustrate the concept behind the measure (see also Supplementary Method section B3). **D** Computational time benchmarking results. Left: Boxplots showing processing times in seconds recorded for each tested method on hand-segmented images less than 200MB. Middle: Scatter plot of the number of processed cells per seconds by Nessys versus the size of input image The input image consisted of the E8.75 sample image which was resized by varying either plane sizes or number of planes while keeping the same resolution. The same input parameters were used in each case. Bottom panel: Illustration of the maximum image size which could be processed by each tested method on a lab workstation with specification given on the left drawing. Note that the largest tested image was 120 GB which was successfully processed by Nessys, thus, the dashed arrow indicates that Nessys can process larger images.

The table of 3D error maps shown in Fig. 3 hinted at the fact that Nessys does perform particularly well in regions where nuclei are densely packed or where the cells are flat and overlapping (monolayer and acini). We did notice a slightly higher proportion of missed nuclei in embryos compared to non-nessys methods. When inspecting individual images, we were able to confirm that this issue could mainly be attributed to the loss of NE signal when the cells undergo mitosis. Overall, this preliminary analysis indicated that NesSys perfoms comparably with other tools for relatively simple tasks (e.g. blastocysts) and may have an advantage in more challenging contexts.

### Quantification of performance metrics of NesSys in comparison with other methods using the DISCEPTS dataset

To more throughoughy assess the performance of the tested methods on the DISCEPTS dataset, we represented precision and recall for each method and biological specimen as a radar plot (Fig. 4A). Precision measures the proportion of the shapes in the output of a method which correspond to actual shapes in the input dataset while recall indicates the proportion of shapes in the input dataset that have been recognised by a method (see also supplementary materials section B 1 and 2). The F-measure, which combines precision and recall, is given in Table S2 together with additional performance measures as described in (Coelho et al, 2009) and Table S3 for reference.

We confirm that all methods perform well for segmenting nuclei in blastocysts, where nuclei are generally near-spherical in the densest region (ICM) or well separated (TE). However NesSys achieved close to or above 95% precision and recall where other methods failed to achieve satisfactory results (Fig. 4A). In particular, the greatest discrepancy was observed for images of cells grown in culture (monolayer and acini) where the cells are often flat, overlapping and very heterogeneous in shape.

The summary of error counts shown in (Fig. 4B) confirms this analysis. Importantly, the balanced proportions of under and over-segmentation events and the minimal proportion of missed or spurious events provided a good indication that parameters were properly adjusted for all methods.

Ultimately, the purpose of automated nuclear segmentation is to fulfil the need for the quantification of biologically relevant features. Therefore, we then assessed the deviation of automated methods from GT in the distribution of generated features related to shape, intensity or neighbourhood (Fig. 4C).

One of the most frequent application of automated segmentation in biology is its use in generating FACS-like profiling of gene expression from immunofluorescence data, i.e quantitative immunofluorescence (Davies et al., 2013; Malaguti et al., 2013; Muñoz Descalzo et al., 2013; Tsakiridis et al., 2014; Wymeersch et al., 2016; Zhou et al., 2013, 1). To test performance on this aspect, we simulated the signal from three heterogeneously expressed transcription factors and computed a distance from GT intensities (see Supplementary Methods B3). We observed that Nessys, Ilastik and MINS performed similarly with a median of the distribution of distance under 20 AFU (3 channels). This was an interesting observation indicating that intensity measurement was relatively robust to segmentation errors, notably MINS error counts was greater in all biological samples and the JI deviation from GT one of the most important (Fig.4C) whereas Nessys’ Jaccaird Index (JI) was the closest from GT.

Nuclear shape features are particularly relevant when studying morphogenesis (Dufour et al., 2017; Etournay et al., 2016; Guirao et al., 2015; Reuille et al., 2015; Stegmaier et al., 2016; Veeman and Reeves, 2015; Xiong et al., 2014). For this reason we computed the shape’s anisotropy and the angle between the main axis of the best fit ellipsoid for a detected shape and its matching GT shape. In this case segmentation errors were much less forgiving, only Nessys and Ilastik provided satisfactory results and when looking at results for individual biological samples, these features deviated from GT for Ilastik where Ilastik segmentation resulted in the greatest number of errors (Monolayer and acini, Fig. S5).

We also assessed performance on the detection of the number of neighbours as this feature can be particularly relevant when studying collective organisation of the cells in various contexts (Blin et al., 2018; Bove et al., 2017; Mesa et al., 2017; Schmitz et al., 2017; Shaya et al., 2017; Toth et al., 2018). Again, only Ilastik and Nessys resulted in an accurate neighbour count with a standard deviation of 2.2 neighbours from GT. This could be explained by the particular ability of Nessys and Ilastik to identify precisely the position of the shapes centroid and to result in shapes with higher JI than other methods (neighbourhood identification uses both centroids and the shape of the nuclei (see Fig. 4C bottom right image and sup methods B.3)

Altogether, our data provide an overview of the level of accuracy that can be expected from Nessys segmentation on biologically relevant features and we conclude that Nessys is a satisfactory method for applications involving measuring intensities, anisotropy, polarity and neighbour analysis.

Finally, we addressed the problem of processing time becoming limiting for large images. Using a desktop computer with 64GB of RAM, no method other than Nessys was able to process images larger than 2GB in our hands, whereas Nessys was able to process a time lapse image dataset of more than 120GB (not shown). This property of Nessys was enabled by the underlying SCIFIO I/O library (Hiner et al., 2016) and ImgLib2 image structure (Pietzsch et al., 2012, 2) which allowed to process each time frame sequentially and which handled lager than memory images. Processing limitations depend on the computing resources available, but Nessys has the advantage of making it feasible to use a typical lab computer to segment large images.

Some of the methods we tested failed to complete segmentation or resulted in prohibitively long segmentation times when images were larger than 200MB, we therefore report processing times for non-Nessys methods for image crops <200MB only (Fig. 4D and Table S4). While in some cases, processing time was shorter than with Nessys, Nessys was on average faster than other methods, and never exceeded a processing time of 132 seconds while maximum processing time was found to be 251, 660 and 702 seconds for Ilastik, Farsight and Mins respectively.

We noticed that with other methods, processing time increased proportionately with image size (Table S5) which was not the case with Nessys. To investigate this aspect and as Nessys was capable of segmenting large images we artificially varied image size of the E8.75 image by either varying the size or the number of image planes. We observed that the time taken to segment each nucleus remained constant regardless of image size with an average of 40 nuclei per seconds for the set of parameters used in this experiment (Fig. 4D). In other words Nessys processing time increases linearly with the number of nuclei in the image and not with image size. This property can be explained by the fact that Nessys only uses 2 image filters which need to work on every voxel of the image and these filters work in parallel on individual 2D planes. Subsequent steps work on detected maxima which quantity is proportional to the number of nuclei in the image.

Overall, we conclude that Nessys performs particularly well for large images and 3D images containing crowded nuclei and provides a useful addition to the toolkit of segmentation approaches currently available.

### Measurement of gene expression and morphological features in post-implantation embryos

To illustrate the usefulness of Nessys segmentation and the associated 3D painter for the study of large embryos, we made use of mouse embryos that we have engineered to express venus under the control of the regulatory elements of Tcf15 (Lin, Tatar et al, in preparation), a transcription factor that marks the formation of somites (Burgess et al., 1996). The signal of the Tcf15 venus reporter is included in the E8.75 image of the DISCEPTS dataset. Using Nessys segmentation on the full image, we generated a plot of the median Tcf15 intensity per nucleus versus position of nuclei along the antero-posterior (AP) axis of the embryo (Fig. 5A). We observed that within the fraction of cells which expressed higher levels of Tcf15 than average (> 1000 AFU) several clusters of points along the AP axis were clearly apparent. Using Nessys 3D painter to manually annotate individual somites as well as other anatomical landmarks (Fig. 5B, D), we confirmed that these point clusters reflected spatial organisation of the somites.

**Fig. 5.**
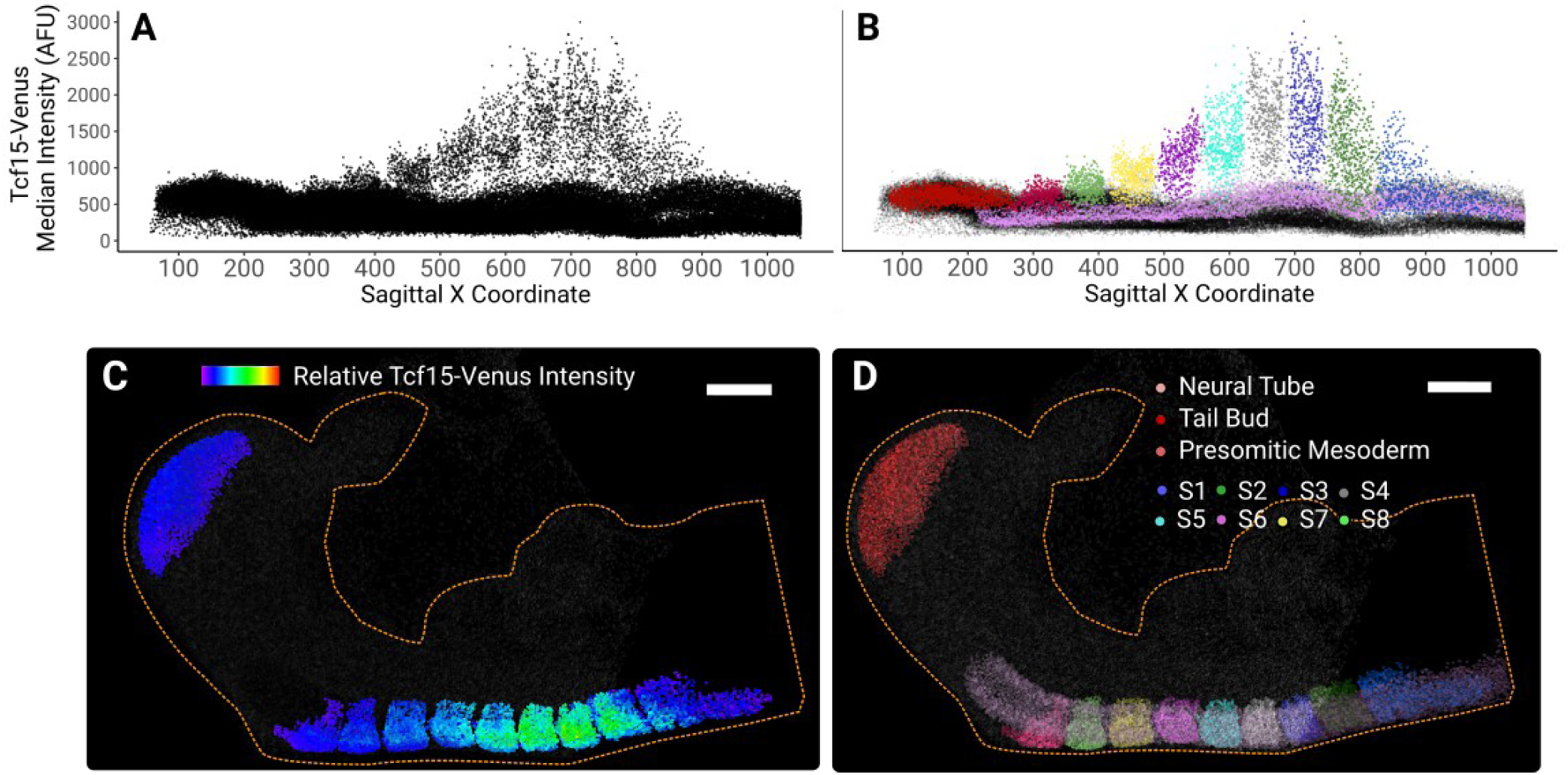
Quantitative analysis of Tcf15 expression in a complex E8.75 mouse embryo. **A** Scatter plot of the median Tcf15-venus intensity within the cell versus the cell position along the Sagittal X coordinate. Notice the groups of points regularly organised along the X axis when Tcf15 is high. **B** Same as A except that points are colour coded according to their respective embryonic region as shown in D. **C** 3D rendering of the all detected nuclei in the E8.75 mouse embryo image. The heatmap of the relative Tcf15-venus intensity is shown only for the tailbud and somitic regions, other regions are shown in grey. **D** 3D rendering of nuclei grouped by embryonic regions. Scale bar in C and D: 100μm, the yellow dashed outline delineates the shape of the embryo.

Tcf15 was known to be expressed in somites but no analysis at single cell resolution has previously been reported. We observe a gradual increase in Tcf15-Venus fluorescence as somites age, reaching a peak at the 7^th^ oldest somite (S3), after which expression declines sharply in the oldest somites S1–2 (Fig. 5A, B, C). Notably, low levels of expression were also detected within the presomitic mesoderm prior to formation of the first somite. We also observe that Tcf15-Venus expression displays considerable heterogeneity within each somite. This analysis illustrates how the combination of Nessys segmentation and 3D annotation tool can be used to quantify large multidimensional images of complex embryos with good accuracy and within a reasonable time frame.

### A fluorescent NE reporter for the automated tracking of live cells

Next, we set out to label the NE in live cells in order to leverage Nessys segmentation power as an input for cell tracking.

While live markers of the whole nucleus are well characterised (for example H2B-FP (Kanda et al., 1998), FP-NLS (Kalderon et al., 1984) and commonly used for cell tracking (Amat et al., 2014; Faure et al., 2016; Lou et al., 2014; McDole et al., 2018; Stegmaier et al., 2016; Ulman et al., 2017; Wolff et al., 2018), this is not the case of NE markers.

An ideal live reporter needs to be bright, accurately localised and non-disruptive to cell function when constitutively expressed. Lamins are well known to regulate cell function and even small changes in their sequence or their expression levels can have deleterious effects (Dittmer and Misteli, 2011; Gruenbaum and Foisner, 2015). Similarly fluorescent tagging of other NE associated protein may disrupt the function of the NE and affect cell function (Crisp et al., 2006). Given these considerations, we generated NE-mKate2, a chimeric construct consisting in the bright mKate2 fluorescent protein (Shcherbo et al., 2009) linked to a NLS and attached to the single-pass trans-membrane domain of Emerin (Bengtsson and Wilson, 2004). We reasoned that this construct would be constantly imported into the nucleus while being tethered to the intra-membranar network of the cells (Fig. 6). Indeed, random integration of NE-mKate2 driven by a CAGS promoter (Alexopoulou et al., 2008) into mES cells resulted in a bright signal with a robust localisation to the nuclear rim (Fig. 6). As this construct is devoid of any known protein interaction motif apart from the NLS, we expect this protein to be inert to cell function. Notably, we confirmed that NE-mKate2 ES cells were able to contribute at high efficiency to chimeric embryos, indicating that the fluorescent label had no obvious deleterious effect on the cells (Malaguti et al in revision). To conclude, the NE-mKate2 construct is a novel non-disruptive fluorescent reporter for detecting the nuclear envelope in live cells.

**Fig. 6.**
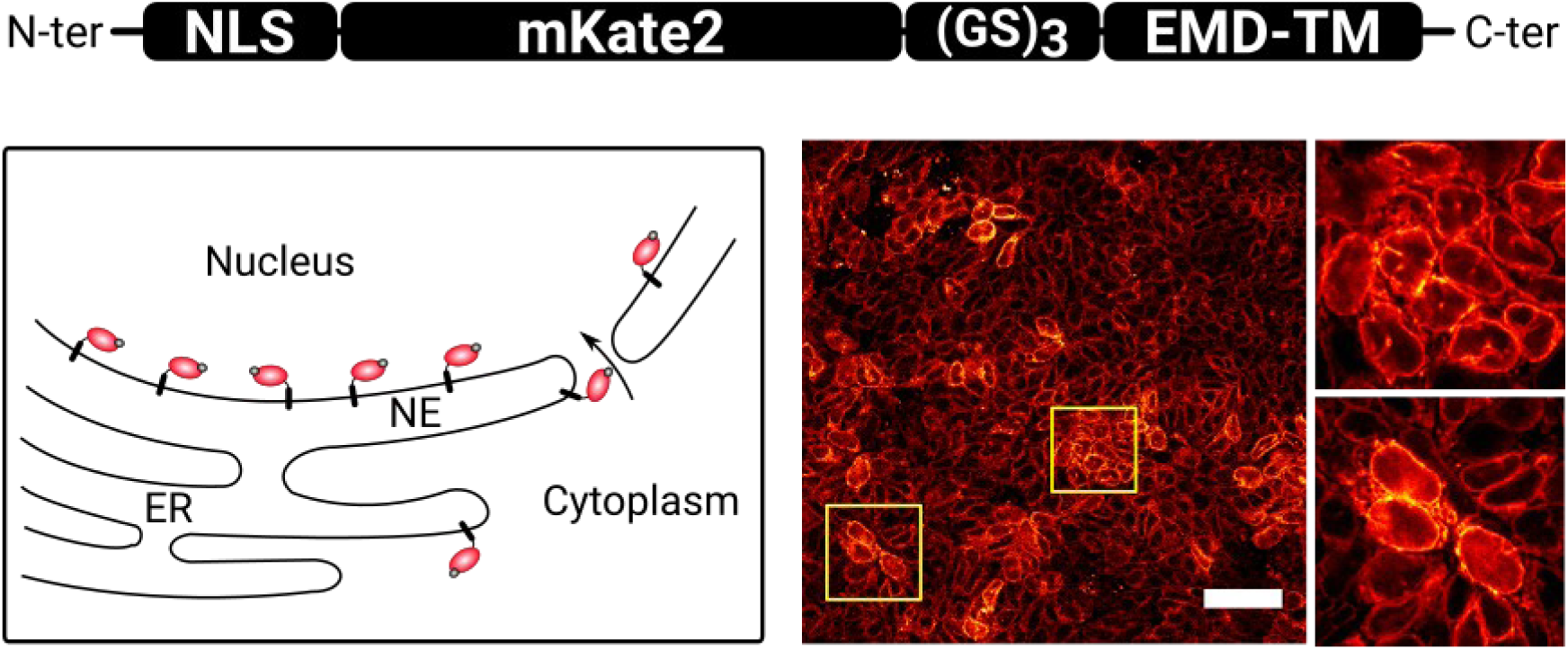
A novel synthetic NE fluorescent protein for live cell tracking. **A** Schematic representing the structure of the NE live reporter and its expected topology and localisation within the cells. NLS: Nuclear Localisation Signal, (GS)_3_: Gly-Ser linker, EMD-TM: 44 C-ter amino acids from the human emerin protein (UniprotKB: P50402) which contain a transmembrane domain **B** High resolution confocal image showing the reporter localisation within a stable cell line constitutively expressing the reporter.

### Tracking cell-cell interactions during differentiation of pluripotent cells

To explore the utility of our tools for identifying cells within time-lapse datasets, we performed confocal live imaging of pluripotent cells differentiating into neural cells. We used mouse ES cells containing a Sox1-GFP reporter (Ying et al., 2003) in order to detect Sox1+ neural progenitors as they emerge over time (Fig. 7A, Movie S3). We chose this system because during the first few days of this process, flat cells form tight 3D clusters in which it becomes difficult to distinguish individual cells, (Fig. 7A). Notably, the neural monolayer culture condition was amongst the most challenging dataset to segment according to our benchmarking results (Fig. 4A and B).

**Fig. 7.**
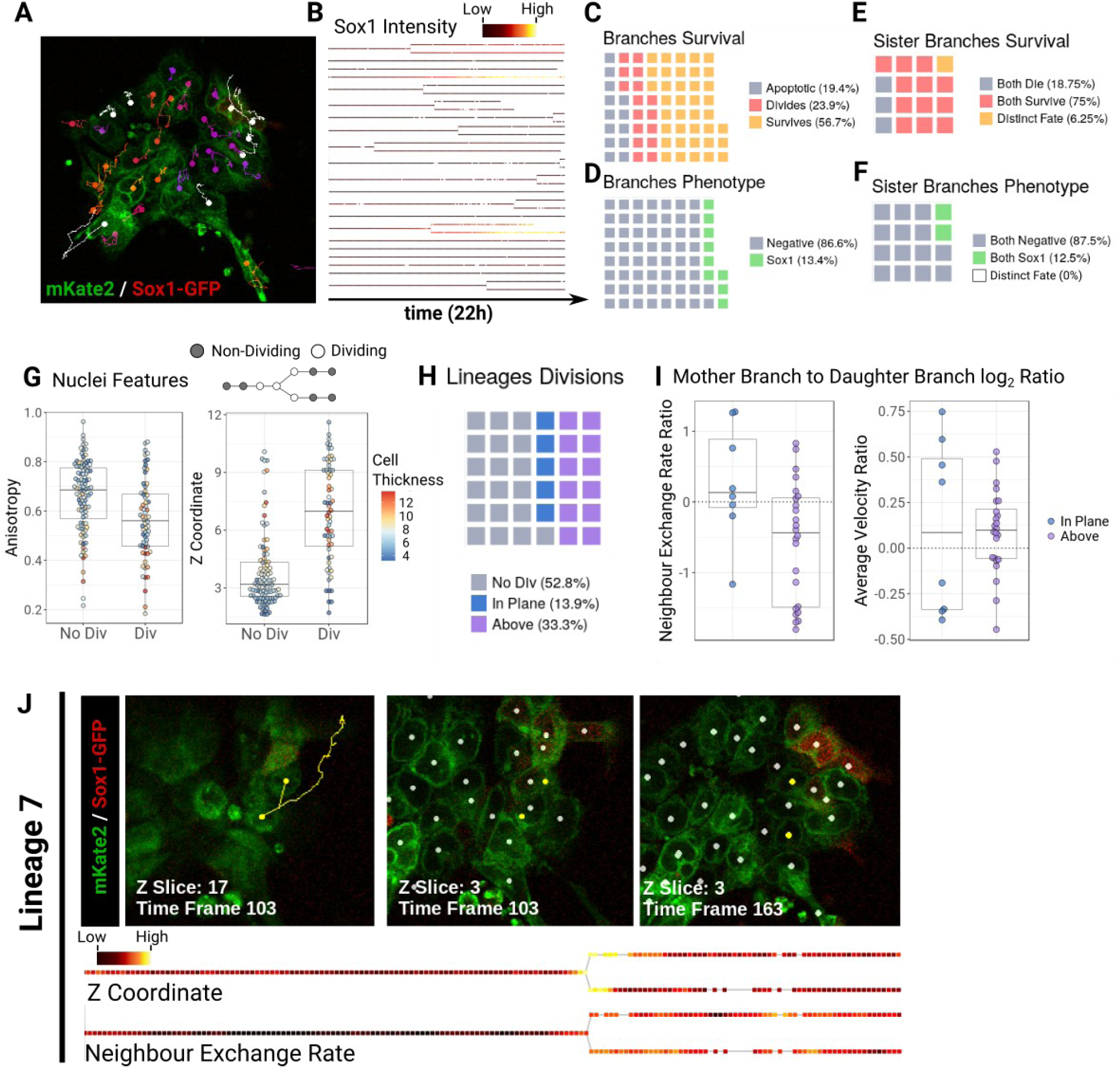
Lineage and neighbour exchange analysis reveal that cell mixing is favoured by divisions above the epithelial plane. **A** Single frame from the Sox1 time lapse experiment. Individual tracks are overlaid on top of the mKate2 (green) and Sox1 (red) signals. The dots indicate the current position of the detected cells. **B** Tree representation of identified lineages. The level of Sox1 intensity within the cells over time is shown with black to red to yellow colour code. **C-D** Waffle charts representing the destiny of individual lineage branches either in terms of apoptosis, survival or division (C) or in therms of Sox1 identity (D). **E-F** Waffle charts comparing the destiny of sister branches for survival (E) or for differentiation (F). **G** Beeswarm-boxplots showing the Z coordinate or the anisotropy of dividing and non-dividing nuclei. A schematic is provided on top of the plots to illustrate which nuclei are considered as ‘dividing’. **H** Waffle chart giving the number of lineage trees contain a division occurring above the epithelial plane, within the plane or which do not contain any division. **I** Beeswarm-boxplots showing the mother branch to daughter branches log_2_ ratio of the average velocity or neighbour exchange rate. **J** Representative track (yellow) containing a division above the epithelial plane which leads to greater neighbour exchange rate after division (white dots represent other detected cells in the image). The corresponding colour-coded trees based on Z coordinate or neighbour exchange rate are shown below images (gaps in the tree correspond to time frames within which the cell was not detected). Note the high Z coordinate at division and the increase in neighbour exchange rate after division.

Although excellent tools exist for the automated live tracking of 2D cultures (Bove et al., 2017; Hilsenbeck et al., 2016; Piccinini et al., 2016; Roccio et al., 2013; Winter et al., 2015), the situation described here would not be possible to address with these tools because of the propensity of differentiating mES cells to squeeze underneath each other (Movie S3) making it necessary to perform analysis in three dimensions.

One possible approach to tackle this challenging situation would be to perform mosaic labelling to make it easier to track a few fluorescent cells among predominantly unlabelled cells. This would be sufficient to extract a subset of cell lineages but would fail to capture local cell-cell interaction or the broader multicellular context surrounding each cell, both of which are central to understanding the emergent properties of differentiating populations (Bove et al., 2017; Toth et al., 2018). For this reason we decided to label all the Sox1-GFP cells with our NE-mKate2 construct in order to analyse the collective behaviours of differentiating cells in 3D.

Unlike snapshot images, where Nessys segmented cells with high accuracy without user intervention, tracking of this challenging time-laspe image required a significant amount of manual correction. However, NesSys segmentation plus manual correction could be completed comfortably within one day (approx 7h) whereas manual segmentation of the entire dataset would have taken around 70h (220 time points with an average of 30 cells per frame and at a rate of 100 cells segmented per hour). After segmenting each individual frame, we ran a basic tracker (Supplementary Methods section C) and were able to obtain accurate lineage trees (Fig. 7A, B, Movie S3).

We first tracked rates of cell death, cell survival, cell division (Fig. 7C) and differentiation (Fig. 7D). Interestingly when we compared the destiny of sister cells, we observed that the majority of sister pairs underwent the same fate in terms of survival (Fig. 7E) and that all underwent the same fate in terms of differentiation (Fig. 7F). This observation is consistent with the idea that differentiation decisions can be made several generations before reporters of cell fate become detectable (Cohen et al., 2010; Costa et al., 2011).

We next monitored incidents of neighbour exchange (Supplementary Methods C), making use of the ability of Nessys and associated tools to track the position of cells in 3D. Neighbour exchange is thought to occur frequently in the mouse epiblast, but it is not known to what extent this process influences the distribution of differentiating cells in culture.

We hypothesised that a major contributor to neighbour exchange would be the position of divisions with respect to the plane of cell growth. Indeed, when we explored the 3D organisation of the cells over time, we noticed that the Z coordinate of the cells increased when cells were dividing (Anisotropy was decreasing slightly as expected, however the change in z coordinate was uncorrelated to the change in cell thickness, Fig. 7G). This raised the possibility that as cells divide above other cells, daughter cells would reintegrate into the colony at different locations thus acquiring new neighbours. To measure this, we set a Z-coordinate threshold above which divisions were considered to be occurring above the plane and we classified lineages accordingly (Fig. 7H). Then, we compared the neighbour exchange rate occurring before division and after division and we observed that when the cells divided above the plane, neighbour exchange rate was more likely to increase, thus supporting our hypothesis. Notably we confirmed that this observation was unlikely to be explained by a change in cell velocity (Fig. 7H right). A division representative of this phenomenon is shown in (Fig. 7K).

While this dataset is too small to draw definitive biological conclusions, it illustrates that in principle it is possible to use Nessys and associated tools to measure the position of each cell in three dimensions and assess cellular behaviours of these cells with respect to their neighbours over time, even in challenging contexts where individal cells are tightly packed and difficult to distinguish by eye.

## Discussion

In this study we set out to explore the idea of using the nuclear envelope (NE) for nuclear segmentation as an alternative to DNA/chromatin labelling. This idea was proposed more than fifteen years ago and preliminary results on small 2D images were promising (Ortiz De Solorzano et al., 2001). Surprisingly however, this idea had not yet, to our knowledge, been revisited for complex 3D datasets. Here we present Nessys, a method tailored to automatically segment nuclei on the basis of the NE. We show that this method performs particularly well for ‘difficult’ datasets, where images contain a large array of nuclear shapes and sizes and where nuclei overlap. It is also a method that is fast and which scales well to large images. We discuss below the strengths and weaknesses of our method and we summarise our effort to make the method easy to adopt by the community and the usefulness of the tools and dataset we provide with this article.

### Features of the NE signal explain Nessys performance

To construct the DISCEPTS dataset we focused in sampling a diversity of nuclei shape, size, texture and 3D tissue organisations. The images we gathered provide a good illustration of the multiple challenges which still need to be addressed in nuclear segmentation and we believe this dataset will be useful as a GT dataset for benchmarking purposes.

Importantly, this dataset illustrates that different cell types harbour different organisations of their chromatin leading to very heterogeneous appearances of the nuclear content. For example, the nuclei of neural cells contain regions of high affinity for dapi which can be seemingly randomly distributed within the nucleus, including at the periphery. This property can confound methods relying on the assumption that brighter regions reside at the centre of the shape. Also, nuclei boundaries which are key features for many methods (Dufour et al., 2017; Kan, 2017; Meijering, 2012; Nketia et al., 2017) become very difficult to distinguish even by eye. These problems are avoided when using a NE staining and this explains in part the success of the Nessys method. Nuclear boundaries remain clear when NE-stained nuclei are in close contact with one another (Fig. 1B) and indeed, our benchmarking results show that the two methods which make use of the NE signal (Nessys and Ilastik) performed particularly well for all embryo images (Fig. 4 A, B, Table S2).

One potential disadvantage of the NE signal is that the interior of the nucleus may become difficult to differentiate from the exterior of the nucleus. As this distinction is easily achievable by the human eye, Nessys uses a naive Bayes classifier trained on a small subset of shapes in order to exclude non relevant shapes (inter-nuclear spaces). This classification ensures that only one channel is required to achieve proper nuclear segmentation and leaves other colours available for other purposes. This contasts with Ilastik, which in our analysis required the dapi channel to perform well.

Our benchmarking results also revealed that the biggest advantage of our methods over the others we tested, Ilastik included, was observed for images of cells grown in culture. Although mammalian embryos are recognised to be amongst the most challenging systems to segment (Amat et al., 2014; McDole et al., 2018; Stegmaier et al., 2016), we show that differentiating stem cells in culture can be equally, if not more difficult to segment accurately as they tend to overlap (Molnar et al., 2016) and be flatter and less regular in shape than their in vivo conterparts. Due to the drop in resolution in the z-axis and in NE signal at the top and bottom of the nuclei, even Ilastik failed to separate overlapping cells (Fig. 4A, B). Nessys performs well as it makes full use of the optimal XY resolution first and then uses a graph colouring strategy to link 2D shapes into 3D volumes.

We would like to emphasise that different methods are appropriate for different contexts, and that our method is intended to complement, rather than replace, existing methods, being particularly well suited for segmenting images with complex 3D arrangements of irregularly shaped nuclei.

### Adoptability of the Nessys method

#### 1) Practicality of using NE over nuclear content

DNA labelling dyes are easy to use and for this reason are widely adopted to mark the nuclei of the cells. Such dyes do not yet exist for the NE and so, usage of Nessys requires that the NE be stained with antibodies. This may potentially impose constraints on applications due to host species restrictions. We used LaminB1 as it is known to be expressed in most cell types with minimal cell-cell variability (Swift et al., 2013), but other NE epitopes could be used: we provide in Table S6 alternative antibodies which provide similar signals in our hands. The use of nanobodies might further improve practicality, reduce antibody penetration issues and be more animal friendly, although we have not tested this approach ourselves (Schumacher et al., 2017). We have also developed a NE live reporter for the use of Nessys for time lapse imaging (Fig. 6 and 7). Given that several simple approaches are available for labelling the nuclear envelope, we anticipate that most users should be able to find a method suitable for their own system.

#### 2) Software usability

The community is in need of open-source, well documented and easy to use softwares in order to accelerate quantitative, reproducible and collaborative research (Cardona and Tomancak, 2012; Carpenter et al., 2013; Meijering et al., 2016). In order to align with this goal, and to facilitate adoptability of Nessys, the software is free and open-source (https://framagit.org/pickcellslab/nessys), the installation process only takes a couple of minutes and does not require particular knowledge in computing. Nessys has also been tested on Linux, Windows and MacOS.

A video tutorial and documentation for Nesys are available online and the user interface includes interactive tools for aiding parameter adjustments (see also the issue tracker which allows comments from the community). The software and associated documentation will continue to be improved in response to user feedback. A longer term goal is to remove the need for parameter adjustments altogether. Notably, we envision the possibility to use a few annotated volumes with the Nessys editor as a training set for automated parameter adjustments.

An important practical aspect of Nessys is that it is designed to be fast and to avoid problems due to processing times-per-cell becoming limiting. We have shown that Nessys can produce accurate segmentation results in a reasonable time frame even for large datasets on a standard lab workstation (Fig. 4)

#### 3) Code portability, extensibility and future work

We developed Nessys using good software design practice. For instance, the user interface uses a modular wizard pattern so that individual subtasks may be improved without modifying others. This will facilitate future improvements of the method. Such improvements may include replacing the last morphological filtering step with a modern shape refining method (Machado et al., 2018) or using more sophisticated classifiers (Domingos, 2012) during the shape ranking process to further improve accuracy. Future work will also focus on enhancing visualisation of segmentation quality in order to speed up the segmentation editing process, for example by including a colour map of the segmentation confidence score as described in (Nketia et al., 2015).

Nessys has initially been developed as a module for our own framework but is designed to be portable to other general purpose frameworks such as ImageJ (Schindelin et al., 2015) or Icy (de Chaumont et al., 2012).

This will help bridge our method with other powerful image analysis components. For example, sing an image restoration method such as CARE (Weigert et al., 2018) upstream of a Nessys segmentation together with downstream sophisticated tracking methods (Ulman et al., 2017; Wolff et al., 2018) will bring us a step closer to the goal of recording and interrogating the emergent properties of collective organisation of the cells in context.

## Conclusion

In conclusion, we hope that the tools presented here will contribute to accelerating quantitative research. We provide a robust 3D nuclear segmentation method together with a 3D editor. The software is easy to install as a stand-alone application and is designed to be portable to general purpose frameworks such as ImageJ. We also provide a dataset and a utility program for segmentation benchmarking in the hope that they will be useful to the community. These tools are free and open-source and designed to work on a standard lab computer with datasets generated from conventional microscopes.

## Methods

### Software Development and availability

NESSys and associated programs are written in java. Our programs only depend on open-source libraries. These include ImgLib2 (Pietzsch et al., 2012) for the Image data structure, SCIFIO (Hiner et al., 2016) and BioFormats (Linkert et al., 2010) for Image Input/Output. We developed our code in Eclipse March 0.3 (https://www.eclipse.org/). Dependencies are managed using Maven, and version control with Git. Source code is available on GitLab at framasoft https://framagit.org/pickcellslab/nessys.

Implementation details, source code and user documentation are subject to changes as we continue to improve the software. Source code and updated documentation may be found on our GitLab repository hosted by framasoft (https://framagit.org/pickcellslab/nessys). We created a git tag ‘v0.1.0’ to label the commit which corresponds to the version of the software that was used for this publication.

### ES cell culture

All mouse embryonic stem cell lines were routinely maintained on gelatinised (Gelatin, Sigma) culture vessels (Corning) in Glasgow Minimum Essential Medium (GMEM, Sigma) supplemented with 10% foetal calf serum (FCS, APS), 100U/ml LIF (produced in-house), 100nM 2-mercaptoethanol (Gibco), 1X non-essential amino acids (Gibco), 2mM L-Glutamine (Gibco), 1mM Sodium Pyruvate (Gibco). For 2i-Lif culture (Wray et al., 2011), the cells were maintained in N2B27 medium suplemented with 3μM of Chiron and 1μM of PD0325911. N2B27 consists of a 1:1 ratio of DMEM/F12 and Neurobasal media supplemented with 0.5% modified N2, 0.5% B27 and 2-β mercaptoethanol (all from Invitrogen). Cell culture was maintained at 37°C with 5 % CO_2_ and routinely tested for mycoplasma contamination.

### Neural monolayer differentiation

Sox1-GFP ES cells (Ying et al., 2003) were first differentiated into EpiSC following the protocol described in (Guo and Smith, 2010) and then further differentiated into neural rosettes as follows: a confluent culture of EpiSC was diluted 1/40 and plated onto growth factor reduced Matrigel (Corning) coated dishes in N2B27 medium containing 10μM of SB435215. The medium was replaced every day and the cells were passaged every 2 days at a ratio of 1/3 - 1/6 onto Matrigel coated dishes until neural rosettes became clearly apparent (5 days). On the last passage, the cells were replated onto 12mm matrigel coated glass coverslips and fixed for immunostaining after 2 days of culture.

### Acini / 3D cell culture

Wild-type E14tg2 alpha ES cells, Tcf15-Het and Tcf15-KO ES cells (Lin, Tatar et al., in preparation) were used for this dataset (The name of the cell lines used for a given image is included in the name of the image). Cells were maintained in 2i-Lif condition and then plated onto Matrigel coated dishes into N2B27/1%KSR medium supplemented with 12ng of bFgf and 20ng of Activin A as described in (Hayashi et al., 2011) and containing 600μg/ml of growth factor reduced Matrigel to induce the formation of 3D structures. The cells were fixed for immunostaining on day 2 of EpiLC induction.

### Mouse strains, staging and husbandry

Wild-type, outbred MF1 mice and transgenic mice - Tcf15-venus (Lin, Tatar et al. In preparation). All mice were maintained on a 12-hr-light/12-hr-dark cycle. For timed matings, noon on the day of finding a vaginal plug was designated as E0.5. Staging of early mouse embryos was done according to (Downs and Davies, 1993; Lawson et al., 2016).

### Embryos collection and dissection

For blastocysts, embryos were obtained at 8 cell morula stage by washing E2.5 oviducts with M2 medium (Sigma). The zona pellucida was removed by a brief wash in Acid Tyrode’s solution (Sigma) at room temperature. Embryos were cultured in KSOM medium (LifeGlobal) at 37C, 5% CO2 for 24-hours prior immunostaining. E7.5 and E8.75 embryos were isolated from the decidua in M2 medium and the Reichert’s membrane was removed before fixation and immunostaining.

### Immuno-fluorescence, embryo clarification and imaging

Samples were fixed in 4% formaldehyde / PBS / 0,5% Triton X-100 for 10 min (cell cultures) or 30 min (embryos) at room temperature. For large embryos, the fixative was quenched with 100mM Glycine/PBS for 5 min. After 3 consecutive washes with PBS/Triton, samples were incubated for a minimum of 30 min (cell culture) or overnight (embryos) with blocking solution which consisted of 5% donkey serum (Sigma), 0.1% Triton X-100 (Sigma) and 0.03% Sodium Azide (Sigma) in PBS. Incubation with antibodies was performed for 1h (cells) or overnight at room temperature (>E7.0 embryos) or at 4°C (blastocysts). Antibodies were all diluted in blocking solution. Antibodies used for each image is given in Table S1 and dilutions and references are listed in Table S6. Samples were counterstained with 1μg/ml Dapi (Sigma) diluted in PBS for 10 min (cells and blastocysts) or 30 min (> E7.0 embryos) at room temperature. Cells grown on glass coverslips were mounted in ProLong Gold Antifade Mountant (Molecular Probes) 24h prior imaging. E7.5 and E8.75 embryos were further clarified using a Benzylalcohol/Benzylbenzoate (BABB) based method adpated from (Dodt et al., 2007). Briefly embryos were dehydrated in graded methanol series (25%, 50%, 75%, 90%, 100% twice) of 5 min each. Embryos were then transferred to a 1:1 solution of BABB/Methanol for 5 min before a final transfer into a pure solution of BABB inside a glass bottom metallic chamber for imaging. Blastocysts were placed on a 10μl PBS drop covered with mineral oil (Sigma) in a glass bottom metallic chamber. Microscope and objectives used for imaging are listed in Table S1.

### Segmentation methods: versions, parameter adjustments and benchmarking

64 bits versions of Ilastik (version 1.1.9), Farsight linux version 0.4.4 and Nessys v0.1.0 were tested on a desktop computer (i7 6700K 8 cores cpu, 64GB of RAM and a 7200rpm HDD) running Linux Opensuse Leap 42.3. MINS version 1.3 was run on the same computer but on a Windows 8 OS with Matlab (version R2016a 9.0.0.341360).

In order to obtain the best segmentation output possible for each method tested, we followed recommendations available for each tool (http://katlab-tools.org/, http://farsight-toolkit.ee.uh.edu/wiki/Nuclear_Segmentation, https://www.ilastik.org/documentation/index.html). For Nessys, Mins and Farsight, we screened 5 segmentation attempts with distinct sets of parameters and retained the output resulting in the best F-measure when compared to the GT (Table S2). For Ilastik, we first trained the pixel classifier on both the LaminB1 and Dapi channels to recognise 3 pixel classes: background, nuclear content and ‘inter-nuclear space’ (z-axis included) which often consisted in a bright LaminB1 signal. Using this method greatly improved Ilastik’s ability to separate touching nuclei in the subsequent steps. We then tested 5 sets of thresholding parameters and retained the best segmentation result as explained above.

Processing times were recorded by starting and stopping a digital stopwatch manually. Nessys also logged processing times to the terminal. Farsight was run from the command line while all the other methods were run from their graphic interface.

Metrics listed in Table S2, summarised in Fig. 4A and B and error maps shown in Fig. 3 were computed with our segmentation comparator program (Fig. S4) and morphometric measurements shown in Fig. 4C and S5 were computed within PickCells (Blin et al. In preparation) as detailed in Supplementary Methods.

### Nuclear envelope reporter construct and cell line

To generate the NE reporter, we used Gibson assembly to ligate a NLS-mKate2 fragment and the human Emerin transmembrane domain (nucleotides 878 - 1012 of NM_000117.2) downstream of a CAGS promoter included in a hygromycin resistance containing backbone. mKate2-NLS was pcr amplified from pTEC20 which was a gift from Lalita Ramakrishnan ((Takaki et al., 2013) Addgene plasmid #30179; http://n2t.net/addgene:30179; RRID:Addgene_30179) and the EMD sequence was pcr amplified from a full length human emerin containing plasmid (kind gift from Dr E. Schirmer). The NLS and (GS)3 linker sequences were included in the primers overhangs.

The resulting plasmid was validated by full sequencing of the insert and will be deposited to addgene.

To generate the Sox1-GFP / mKate-NE double reporter cell line, we lipofected 2.5μg of plasmid (Lipofectamin 3000, Thermofischer) according to the manufacturer’s recommendation into the Sox1-GFP cell line (Ying et al., 2003). Hygromycin resistant clones were selected for the best compromise between high mKate2 intensity and proper NE localisation of the transgene.

### Time lapse Imaging and tracking

For the tracking experiment, the sox1-gfp/NE-mKate2 cell line was maintained in 2i/Lif condition and then differentiated into the neural lineage as follows: The cells were dissociated with accutase (invitrogen) and replated at a density of 10 000 cells / cm^2^ into N2B27/1%KSR medium. We determined that Sox1-GFP became expressed between day 2 and day 3 of differentiation and so replated the cells at day 2 of differentiation into phenol-free N2B27/1%KSR medium inside a glass-bottom metallic chamber for imaging. Imaging was performed with an inverted Leica Sp8 TCS microscope using temperature and CO2 (5%) control, a 20x objective with NA=0,7 and glycerol immersion. Excitation lasers were 488nm and 561nm and signal was captured with HyD detectors. 25 z-planes were captured every 6 minutes, final image resolution was 0.4 × 0.4 × 0.57 μm. The tracking and analysis procedure is described in Supplementary Methods C2.

## Acknowledgements

We would like to thank Dr Chia-Yi Lin for the Tcf15 embryos and Dr Anna Kreshuk for advice in using Ilastik and feedback on segmentation outputs. We are also grateful to all our colleagues who have taken the time to test Nessys and have given precious feedback. This manuscript has been generated with free and open-source software (Zotero, Inkscape, LibreOffice, Linux).

## SUPPLEMENTARY METHODS

### A. Detailed description of the segmentation method

An overview of the segmentation procedure is depicted in Fig. 2. This section provides a more detailed description of some of the key steps included in the segmentation method. Please also note that source code and the latest documentation (including advice on how to adjust parameters) can be found on the Nessys online repository: https://framagit.org/pickcellslab/nessys

#### 1. Tree-structured ridge following procedure (Fig. S1)

The purpose of this step is to identify outlines of nuclei within a given 2D image plane.

Inputs received from the previous steps include:

- The original NE signal for a specific plane
- The corresponding ridge-enhanced plane (steerable filters output)
- An image location to use for initialisation

The procedure starts from a unique location in the 2D plane (maximum identified in the previous step) and follows the ridges in the vicinity. The key idea is that ridges are built incrementally to form a series of ‘ridglets’ organised as a tree until some stop conditions are met. The advantages are several:

- A shape can be constructed from any leaf pair by following the ridges leading to their least common ancestor (LCA). (In the end a classifier ranks the shapes and identifies the leaf pair to be selected).
- As a consequence, ridglets at the top of the hierarchy have a higher probability to be selected as part of a nucleus outline.
- When building a shape from 2 leaves, ridge features can be computed from the sum of pre-computed features for each ridglet to speed up the process.

Fig. S1 depicts this procedure. And Movie S2 shows a real example of the ridge following procedure in slow motion.

#### 2. Depth linkage procedure

Once all planes have been segmented, individual areas are linked together into 3D volumes (Fig. S3). The procedure is as follows:

- Create a directed graph (V, E) of potential connections where V are areas and E define potential linkage (Fig. S3 A, B, C)

- Create an edge E to the shape in the next plane if:

- inter-centroid distance is less than threshold
- shape overlap is more than the minimum percentage of each area
- Create an edge E to the shape in the subsequent planes if

- the number of ‘jumped’ planes is less than threshold
- the shape is not already linked to other areas in previous planes
- Assign edge features for edge ranking during the graph colouring step

- inter-centroid distance
- jaccard index
- sum of overlaps
- centre of overlaps
- Find areas with ambiguity (which mean more than 1 incoming or outgoing edge)

- if multiple connections above and below then split (Fig. S3D). Splitting involves checking which of the above or below merged areas have higher JI and then replace current area with copies of the chosen areas. NB: to account for this later on, the newly created edges will be assigned the JI computed between the merged area that were copied and the area that was split.
- if ambiguity above only, then decide between merging shapes above or splitting the current area using Jaccard indices
- if ambiguity below only, then decide between merging shapes below or splitting the current area using Jaccard indices
- Graph colouring (Fig. S3E)

- Create a sorted queue of edges in descending value of Jaccard index
- For each edge check the source and target area

- Case 1: None have a colour: Create a new unique colour and assign to both
- Case 2: Only 1 has a colour: Test if volume is within bounds and merge
- Merge small volumes at boundaries if merged volume < Mean volume +/-3 sigma
- Post-processing - All are optional (Fig. S3F)

- Split based on volume ‘anomalies’

- Intensity increase
- barycenter displacement
- Merge extrema?
- Remove volumes made of only one plane
- Morphological smoothing (Closing then Opening then add tips - dilation in z with offset)

NB: This procedure is subject to change as we may further improve the method over time. We have setup an issue tracker (https://framagit.org/pickcellslab/nessys/issues) in our GitLab repository which will enable anyone to look at scheduled plans to improve Nessys and also to propose changes and fixes.

### Complementary details for the performance assessment procedure

#### 1 Error counting

The steps outlined below describe the procedure implemented in the segmentation comparator module to identify segmentation errors and accurate detections:

1. For each shape in the GT image, we associate the shape in the tested segmentation image which possesses the biggest overlap. A reference of each matching pair is kept in memory as we iterate over GT shapes(forward match).
2. The same is done in the opposite direction: for each shape in the tested image, we identify the best matching shape in the GT image. The reference for each matching pair is kept in memory (backward match)
3. Shapes are then classified as follows:

a. Miss: the GT shape shares less than 5% of overlap with its forward match (95% of the shape is background in the test image)
b. Spurious: the tested shape shares less than 5% of overlap with its backward match (95% of the shape is background in the GT image)
c. Merge: There is more than one forward match for the given GT shape in the tested image
d. Split: There is more than one backward match for a given tested shape in the GT image.
e. Accurate: There is exactly one backward match and one forward match for a pair of GT/tested shape.

**NB:** Shapes are excluded from the analysis if the matching shape in the GT image is in contact with any of the 6 image borders.

For each GT/Tested image pair given as input to the program, an image with the same dimensionality of the input image is generated. This output image represents a map of the shape classes defined above. GT and detected nuclei are drawn as an outline and given a label which indicates their class as follows: ACCURATE = 1, EXCLUDED = 2, MISS = 3, SPURIOUS = 4, SPLIT = 5, MERGED = 6. Applying a look up table to this image such as the ‘16-colours’ Lut in imagej allows to visualise the accuracy of the segmentation method (Fig. S4 C, see also Fig. 3)

#### 2. Precision and recall

Precision and recall were computed in LibreOffice calc. based on the output of the segmentation comparator.

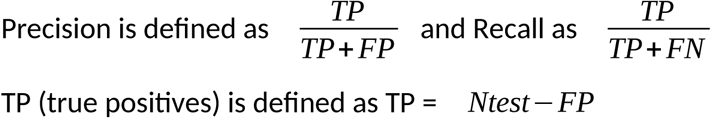

where *Ntest* is the total number of shapes in the test image minus the number of shapes excluded from the analysis and FP (false positives) 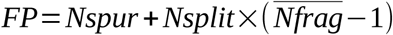 where *Nspur* is the number of spurious shapes, *Nsplit* is the number of split event and Nfrag is the average number of fragments per splitted shape.

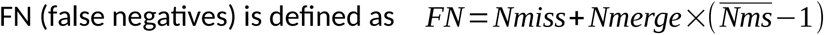

where *Nmiss* is the number of miss event, *Nmerge*, the number of undersegmentation event and *Nms* the average number of shapes per merge event.

#### 3. Morphometric analysis

We performed the computation of morphometric features in a software built in our lab called PickCells (Blin et al., in preparation, source code is available at https://framagit.org/pickcellslab) and exported features to R (R Core Team, 2013) in order to build the plots shown in Fig. 4C and Fig. S5. Features were computed as follows:

##### ■ Anisotropy

To measure the anisotropy for a given 3D shape, we computed the covariance matrix from the list of voxel coordinates of the shape and performed its eigen-decomposition. Our code uses the Math3 Apache library (http://commons.apache.org/proper/commons-math/). Anisotropy was defined as: 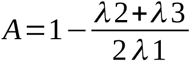 where lambda 1, 2 and 3 are the 1^st^, 2^nd^ and 3^rd^ eigenvalues of the decomposition respectively. The ‘anisotropy difference’ shown in Fig. 4C is the anisotropy value of the tested shape minus the anisotropy value of the matching GT shape.

##### ■ Eigen vector angles

We computed 3D eigen vectors from the eigendecomposition of the covariance matrix of the list of 3D coordinates. The eigen vector angle shown in Fig. 4C is the angle between the first eigen vector of the tested shape and the first eigenvector of the GT shape expressed between 0 and 90°.

##### ■ Jaccard Index

The Jaccard Index (JI) between a tested shape and its matching GT shape was computed by expressing volumes (total volume and volume of the shapes intersection) as a number of voxels. JI was defined as 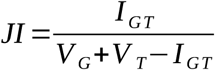 where *I_GT_* is the volume of the intersection between the GT and the tested shape, *V_G_* is the volume of the GT shape and *V_T_* is the volume of the tested shape.

##### ■ Intensity distance

To simulate a heterogeneous expression of 3 transcription factors, we used the 32bit manually segmented output and converted the image to an RGB image and added noise and blur. To compute the intensity distance, we then measured the average intensity for each RGB channel to obtain a 3D ‘colour coordinate’ for each shape. The ‘intensity distance’ shown in Fig. 4C is the euclidian distance between the 3D colour coordinates of the tested shape and its best matching GT shape.

##### ■ Inter-centroid distance

The inter-centroid distance corresponds to the euclidian distance between the centroid of the GT shape and the best matching tested shape.

##### ■ Relative volume difference

The relative volume difference was defined as *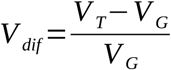*

##### ■ Neighbours difference

To identify neighbours for each individual shape in an image, we first created a Delaunay triangulation using the centroids of the shapes as input. We then removed edges in the graph by applying a cutoff of 5μm to the shortest border to border distance between adjacent shapes. The number of neighbours for a given shape was given by the degree at the corresponding vertex in the resulting graph. The neighbours difference reported in Fig. 4C and Fig. S5 was equal to the number of neighbour of the tested shape minus the number of neighbour of the GT shape.

### C. Additional analysis of Nessys outputs

#### 1. Tcf15 expression analysis

The Tcf15 expression analysis shown in Fig. 5 was performed in PickCells. The Nessys segmentation module was used to perform the segmentation of the full E8.75 image of the DISCEPTS dataset. For Fig. 5C and D, embryonic regions were manually annotated using the Nessys 3D painter module. A rule was created in PickCells to assign nuclei to a given embryonic region if their shape overlapped with the annotated region by at least 95%. Fig 5A and B 3D representations were created using the 3D scene module in PickCells.

#### 2. Cell tracking and Neighbour exchange analysis

The tracking results presented in Fig. 7 were obtained with the tracking module of PickCells (https://framagit.org/pickcellslab/pickcells-essentials/tree/develop/pickcells-tracking) using manually edited Nessys segmentation as input.

#### 1. Neighbour exchange rate definition

First, the neighbour graph was computed as described in (section B3 ‘Neighbour Difference’) for each time point. Then, we compared each nucleus to itself in the previous time point to obtain the number of unique neighbours that have been lost (N_loss_ = set difference of neighbours at t and neighbours at t+1) and the number of unique neighbours that have been gained (N_gain_ = set difference of neighbours at t+1 and neighbours at t) during one time frame:

We then defined the neighbour exchange rate (NER) for a given branch as follows:

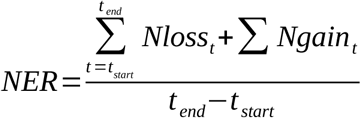

where t_start_ and t_end_ are the time frame of the first and last node of the branch respectively.

#### 2. Nuclei and Branches classification

##### Sox1 classification

We defined a branch as Sox1+ if the mean of average Sox1 intensity of nuclei composing the branch was above a given threshold. This threshold was set subjectively based on both the shape of the distribution of nuclei mean intensities and based on visual inspection of the movie (Movie S3 - PickCells allows us to click on individual nuclei or branches to obtain the features values of the selected object).

##### Dividing nuclei

Nuclei were defined as ‘dividing’ based on their position in the lineage tree, i.e a nucleus with two outgoing links and its depth one neighbouring nuclei in the tree were defined as ‘dividing’ (Fig. 7G).

##### Apoptotic nuclei

apoptotic nuclei were defined as the leaf nodes in a tree which did not reach the end of the movie.

##### Dividing branch

A branch for which the last node corresponds to a dividing nucleus.

##### Apoptotic branch

A branch for which the last node corresponds to an apoptotic nucleus.

##### Surviving Branch

A branch for which the last node was neither a dividing nor an apoptotic nucleus.

##### Above/ below plane classification

We defined a lineage as dividing above the epithelial plane if the average Z coordinates of dividing nuclei was above 6.5 μm which corresponded to the 3^rd^ quartile plus 1.5 times the inter-quartile range of the distribution of Z coordinates of non dividing nuclei (Fig. 7G). Branches were then defined as above or below based on the category of the tree they belonged to. Lineages with no divisions were excluded.

## SUPPLEMENTARY FIGURES AND TABLES

**Fig. S1.**
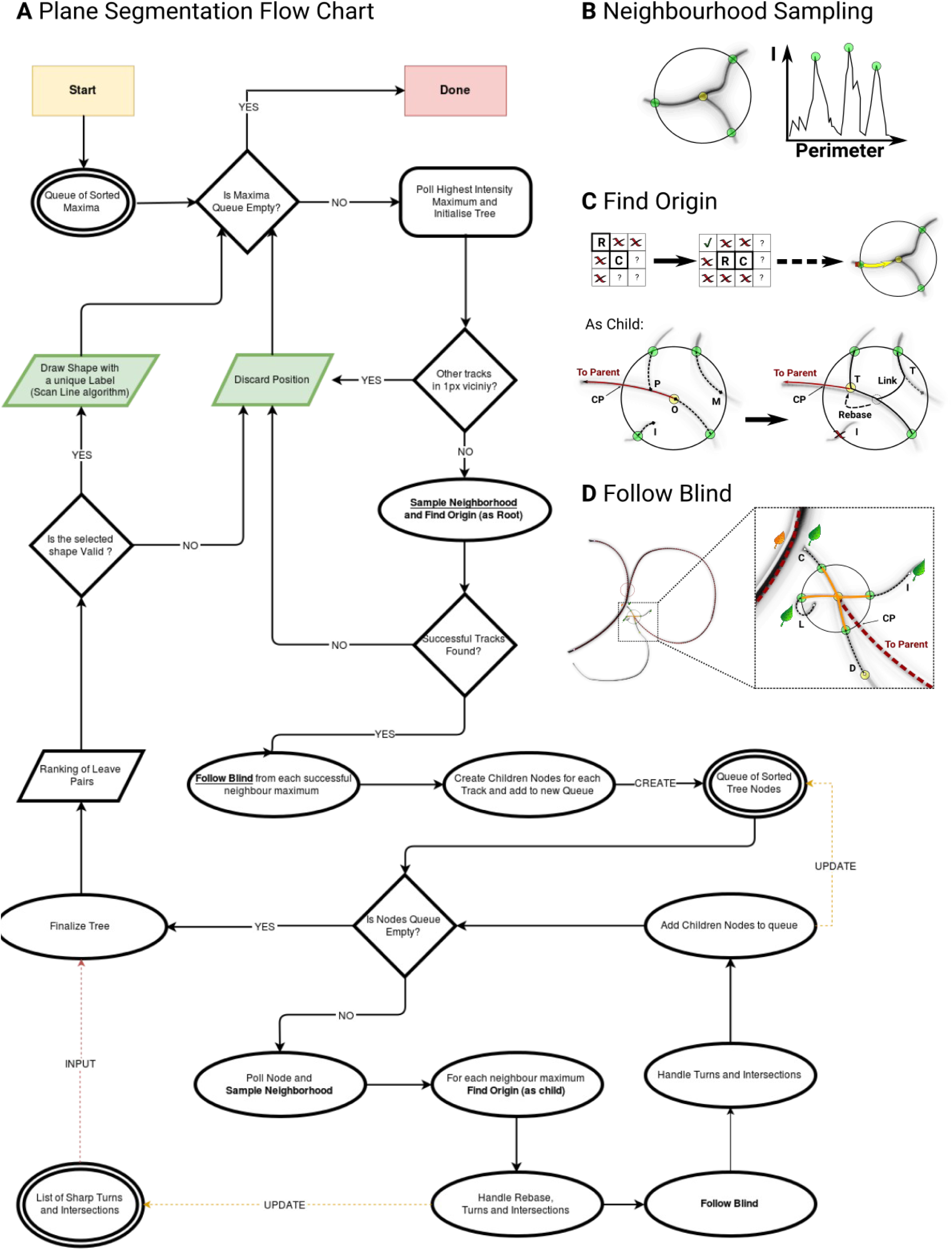
Detailed description of the Tree Structured Ridge Following procedure. **A** Block diagram depicting the flow of operations performed during the ridge following procedure. Double Ellipses: Input/Cache, Simple Ellipses: Subprocesses, Losanges: decisions, Parallelograms: Outputs. Processes in bold and underlined are further described on the right hand side of the flowchart. **B** Describes how the neighbourhood of a node centre is probed in order to follow local ridges. The graph on the right represents the profile of intensities along the perimeter of the circle drawn on the left. **C** Shows how a ridge section is constructed pixel wise. ‘C’ is the current centre of the 8 neighbours window (last pixel identified as belonging to the ridge), ‘R’ represents the pixel identified as ridge just before ‘C’ and is used to find the 3 next potential ridge pixels which are shown with a ‘?’. The schematic on the right shows the direction of the following process from the neighbour maximum towards the node center (Origin). Schematics in the bottom section depicts the stop conditions (I, O, M, P) and the resulting paths after required operations such as ‘rebase’ or ‘linkage’ operations. Note that when this step if performed “as root”, only I and O are used. **D** Illustrates the last step required to construct the ridges to the next nodes in the tree (‘Find Blind’). A large field of view with 3 nuclei is drawn on the left. Branches of the tree that have already been built in this example are represented with red dashes, a red circle is drawn at the position of the root node. The region where the ‘find blind’ step is being performed is magnified on the right. Each possible stop condition is depicted. Green leaves are represented where leaf nodes are identified and an orange leaf is drawn where a path intersection is found as this induces a new leaf child in the corresponding internal node of the tree. Yellow Circle: Current Node Center, Green Circle: Neighbourhood Maximum, CP: Crossing Point, P: Path to parent node, O: Origin, M: Missed, I: Intensity Drop, T: Sharp Turns, C: Own path Crossing. L: Loop to self, Distance stop.

**Fig. S2.**
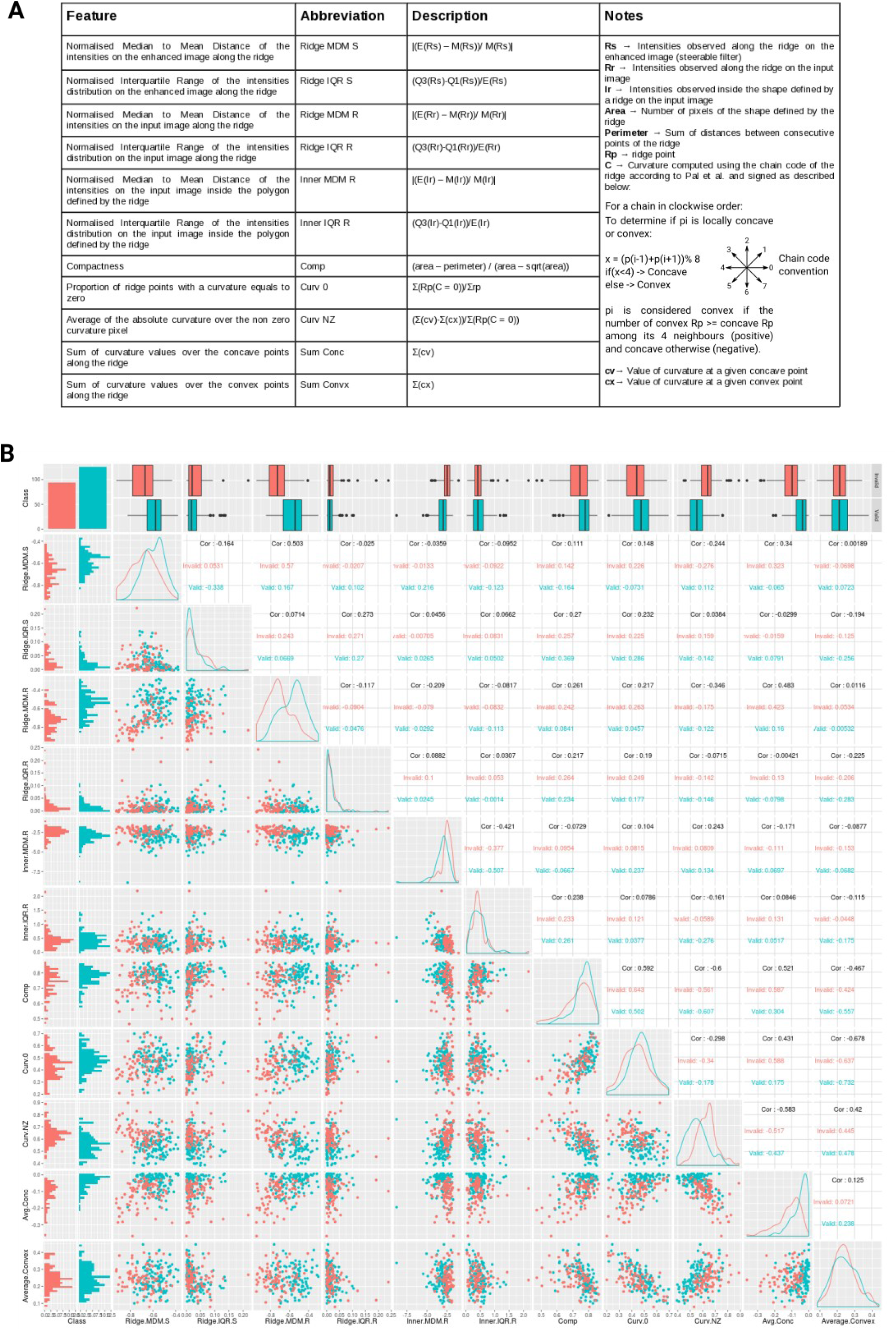
Description and example of distributions of the ridge features used as input to the Naive Bayes Classifier. **A** This table provides an overview of the features computed for each ridge passing the minimal validity check in order to rank and select the most probable shape with a naive Bayes classifier. **B** A matrix of plots representing the distributions of feature values of a sample training set used by the classifier. Blue are valid shapes and red invalid shapes. Plot created in R with ggpairs from the GGally library(“Extension to ‘ggplot2’ [R package GGally version 1.2.0],” n.d.)

**Fig. S3.**
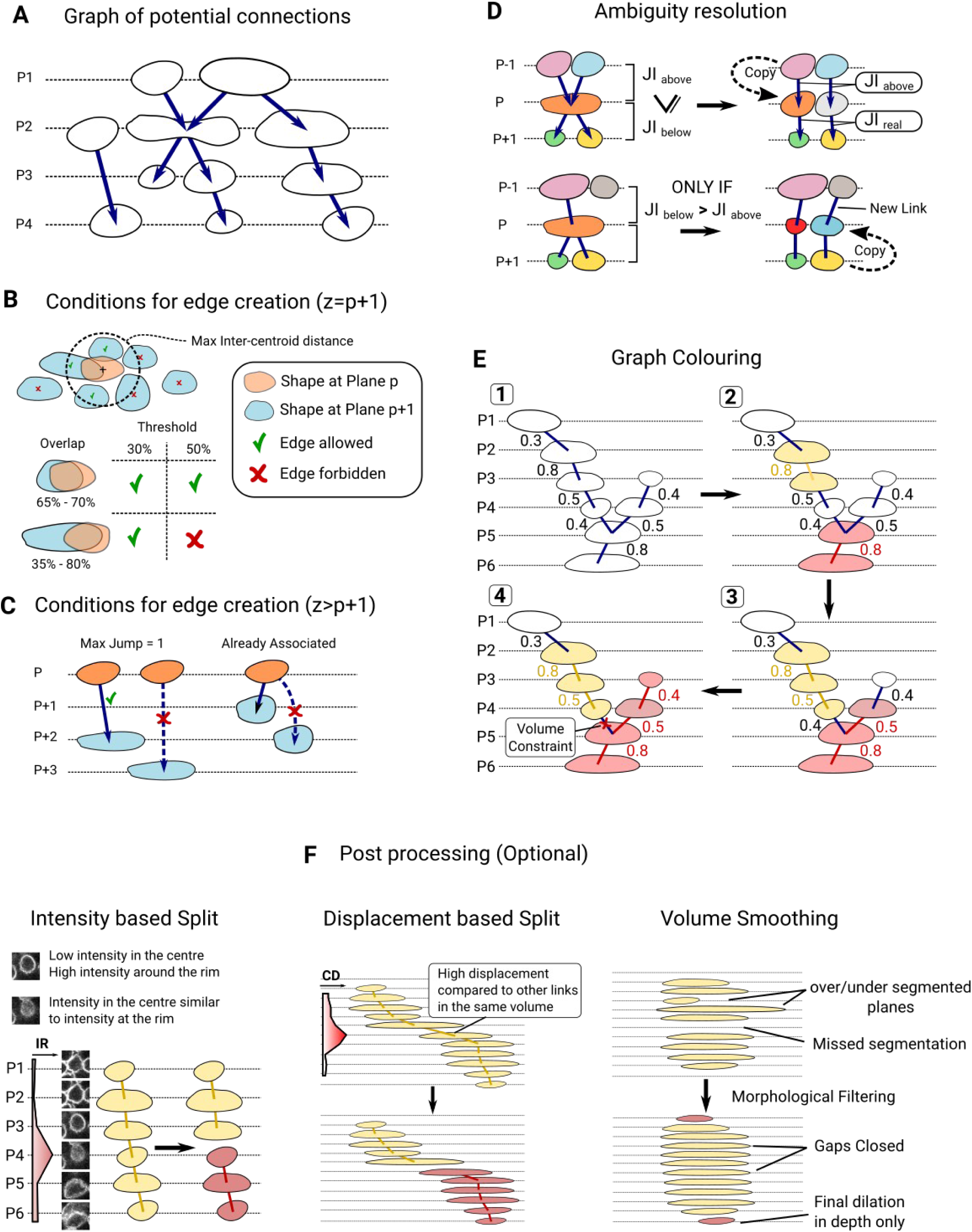
Description of the Depth linkage procedure and definition of user-defined parameters. The schematics in this figure represent individually segmented planes in a 3D image. image planes are shown as dashed vertical lines with 2D areas identified in the previous steps of the segmentation drawn as solid shapes. The relative position of each plane is indicated by the letter P followed by the plane index. **A** Represents an example of a directional graph created by the first step of the procedure. Arrows represent the links which are created if the two connected shapes can potentially be part of the same volume. **B** Required conditions for link creation when the areas to lookup (blue shapes) are located in the directly adjacent plane as the area under consideration (orange shape). The drawing at the top shows the max inter-centroid distance threshold as a dashed circle. The red cross shows the centroids that are excluded and green ticks the centroid that can be included. The bottom drawing shows the effect of the ‘overlap threshold’. Percentages indicate how much of the area overlaps with the surface of the other area. Left percentage is for the blue shape and right percentage is for the orange shape. The outcome with two distinct values for the overlap threshold is given. **C** Condition for edge creation when the area to lookup (blue shape) is located further than the plane directly adjacent to the area under consideration (orange shape). The ‘Max Jump’ parameter defines the maximum number of planes that are allowed between two areas for an edge to be created. The drawing also shows the rule that applies when the area already possesses an edge with an area in a plane located upstream. **D** Diagrams showing rules applying when ‘ambiguities’ are detected. The shapes colours indicate their unique id and the area with ambiguities is shown in orange. Two cases are represented: 1) ambiguity is found both with the above plane and with the plane below. In this example, the Jaccard Index (JI) between the merged areas above and the current area is higher than the JI between the merged areas below and the current plane 2) ambiguity is found only in the plane below. The outcome obtained in the case that the JI between the current area and the merged areas in the bottom plane is higher than the JI between the current area and the area above is shown on the right. **E** Illustration of the volume labeling procedure. Numbers in rounded squares indicate the order of events. The numbers next to the edges represent the value of the JI between two connected areas. Each colour indicates a unique id. Areas with the same id will belong to the same volume eventually. White means that no volume id has yet been assigned. The effect of the ‘volume constraint’ (Max volume and min volume parameters) is shown. **F** Illustrations of the optional post-processing steps. Left panel: Intensity based splitting. Example image planes with Lamin B1 signal are shown. The corresponding diagram illustrates a cut where the peak in the intensity ratio (IR - intensity at the center dividied by intensity at the rim) is found. Middle panel: Displacement based splitting: The top diagram shows an example of an under-segmented volume. The profile of the centroid displacement (CD) along image depth is given. In the bottom drawing, a cut is shown at the peak of the CD value. Right Panel: Volume smoothing. Examples of artifacts are given in the top diagram. The bottom drawing shows how these artifacts are expected to be corrected. Additional areas at the tips of the volume are also shown.

**Fig. S4.**
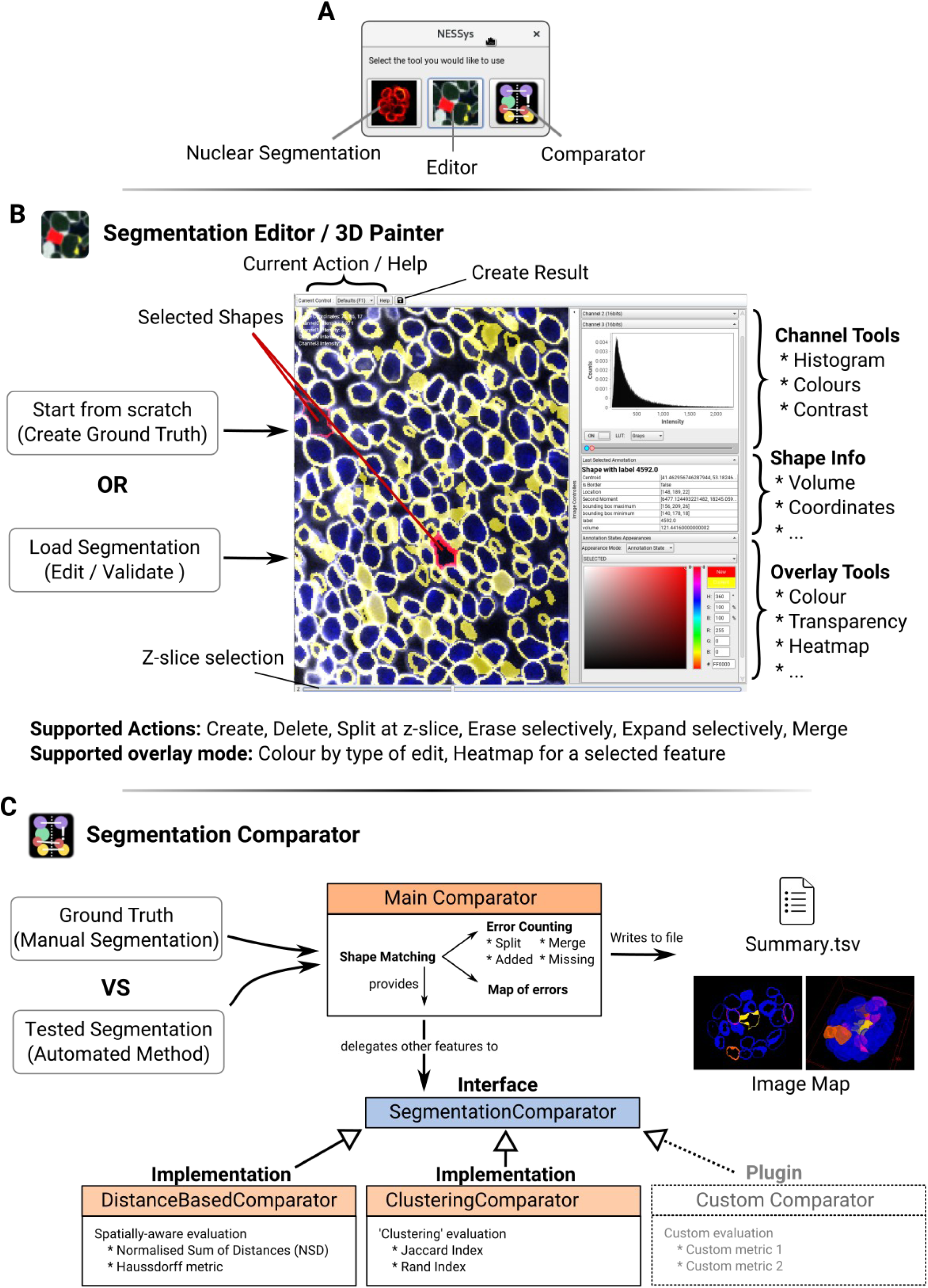
Overview of Nessys features for segmentation validation and benchmarking. **A** Snapshot of the first window to appear when launching the Nessys standalone application. This window lets the user choose the specific tool to work with. **B** Snapshot of the segmentation editor interface. The streak region of the E8.75 embryo (blue and white image) is loaded into the editor together with a segmentation result which appears as yellow outlines. Selected shapes have been highlighted in red. **C** Overview of the ‘segmentation comparator’ tool. Boxes with a title over an orange background represent java implementations in the application. The tasks that they handle is indicated inside the box. The ‘SegmentationComparator’ interface is shown in a blue box and can be extended to add custom metrics to the benchmarking process (dashed box). Required inputs are shown on the left-hand side: A GT image and a tested segmentation are loaded into the application which performs ‘shape matching’, error counting and computes performance metrics. Note that batch processing of images is supported as long as the number and dimensions of GT images are the same as tested segmentation images. Outputs of the program are illustrated on the right-hand side of the diagram. Summary.tsv is a table with all computed metrics for each image. The raw image of the 3D error map created by the program is shown with the ‘16 colours’ lookup table of ImageJ (Image Map - left) and a 3D view of this image obtained with the ImageJ 3D viewer (Image Map - right). Blue: accurate hit, yellow: spurious, purple: merge, orange: split. An online video tutorial for the Nessys method is readily available at https://peertube.mastodon.host/videos/watch/ca891fb7-df04–4da1-abf9–509586162944 and detailed tutorials for all Nessys tools will be released soon on the following website: https://pickcellslab.frama.io/docs/

**Fig. S5.**
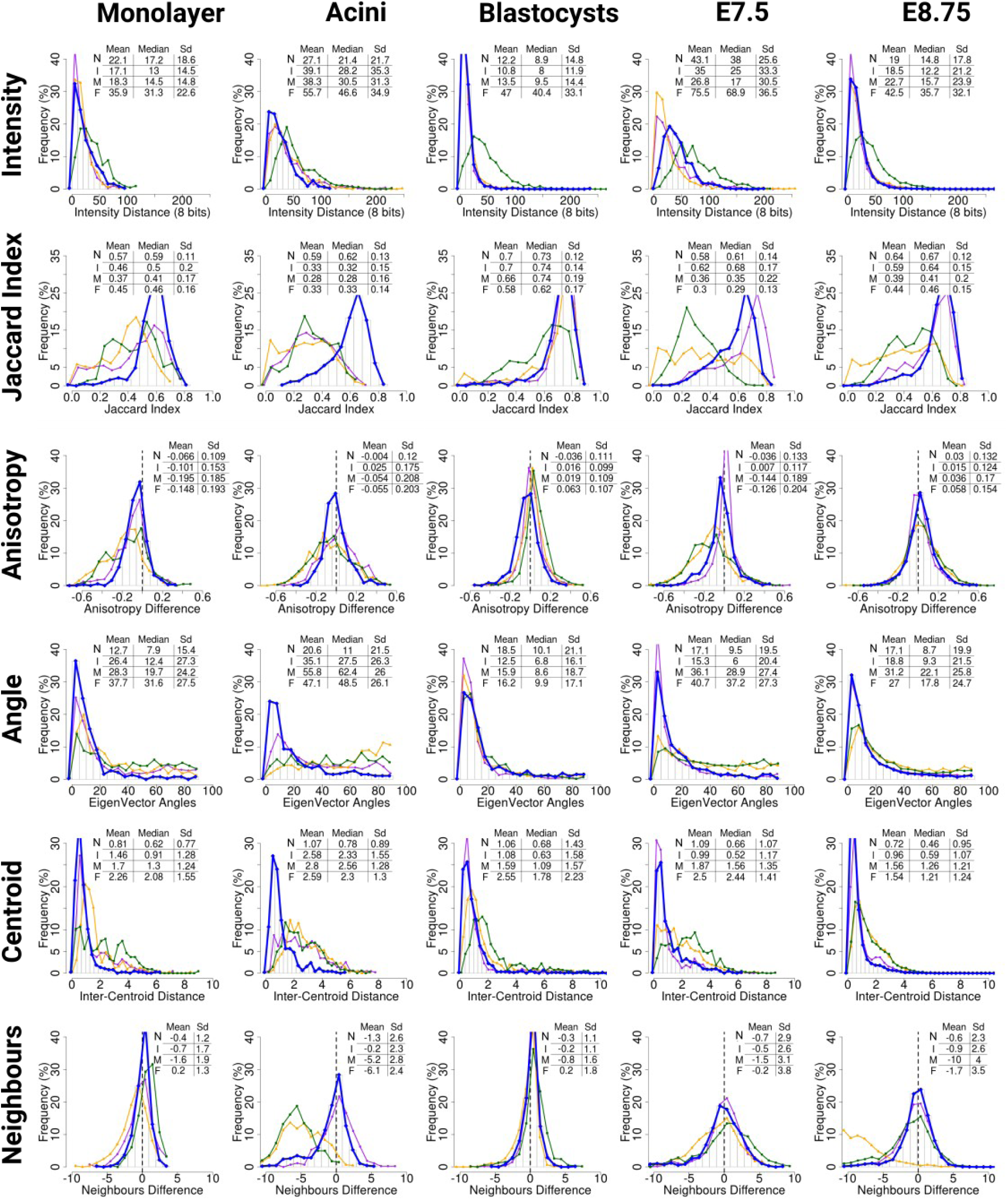
Histograms of the deviation of morphological features from the ground truth for each biological dataset including all nuclei (complementary data for Fig. 4C). blue: Nessys, purple: Ilastik, yellow: Mins, green: Farsight.

**Table S1:** Summary of DISCEPTS image properties.

| Biological Specimen | Number of images | Bit Depth | Channels | Imaging medium | Microscope | Objective | Voxel size (µm) |
| --- | --- | --- | --- | --- | --- | --- | --- |
| Neural Monolayer | 1 | 8 | GammaTubulin / LaminB1 / Sox1-GFP / Dapi | Prolong Gold | Leica SpE | ACS APO 63.0x1.30 Oil | 0,227 x 0,227 x 0,503 |
| 3D Acini | 3 | 12 | Oct6 / LaminB1 / Dapi / gammaTubulin | Prolong Gold | Leica Sp8 | HC PL APO 40x/1.30 Oil CS2 | 0.238 x 0.238 x 0.5 |
| Blastocysts | 5 | 12 | LaminB1 / Dapi | PBS | Leica Sp8 | HC PL APO 40x/1.30 Oil CS2 | 0.36 x 0.36 x 0.5 |
| E7.5 | 1 | 12 | LaminB1 / Dapi | BABB | Leica Sp8 | HC PL APO 40x/1.30 Oil CS2 | 0.142 x 0.142 x 0.5 |
| E8.75 | 1 | 12 | Dapi / Tcf15-venus / gammaTubulin / LaminB1 | BABB | Leica Sp8 | HC PL APO 40x/1.30 Oil CS2 | 0.28 x 0.28 x 1 |

**Table S2:** Segmentation accuracy measures for each biological specimen and method.

| Monolayer ( 1 image - 470 Cells) |  |  |  |  |  |  |  |  |  |  |  |  |
| --- | --- | --- | --- | --- | --- | --- | --- | --- | --- | --- | --- | --- |
| Method | Jl | RI | Hausdorff | NSD | Merge | Miss | Split | Spurious | Accurate | Precision | Recall | F-Measure |
| Nessys | 0,61 | 0,94 | 256,70 | 0,30 | 4,50 % | 2,00 % | 2,84 % | 1,03 % | 89,63 % | 96,13 % | 93,48 % | 94,79 % |
| Ilastik | 0,58 | 0,94 | 261,47 | 0,44 | 10,00 % | 4,25 % | 19,69 % | 2,21 % | 63,85 % | 78,10 % | 86,10 % | 81,90 % |
| MINS | 0,42 | 0,91 | 256,36 | 0,64 | 16,75 % | 3,25 % | 9,25 % | 8,75 % | 62,00 % | 82,00 % | 80,39 % | 81,19 % |
| FarSight | 0,63 | 0,93 | 256,11 | 0,18 | 24,25 % | 0,00 % | 21,32 % | 0,76 % | 53,67 % | 77,92 % | 75,99 % | 76,94 % |

| Acini (3 images - 512 Cells) |  |  |  |  |  |  |  |  |  |  |  |  |
| --- | --- | --- | --- | --- | --- | --- | --- | --- | --- | --- | --- | --- |
| Method | Jl | RI | Hausdorff | NSD | Merge | Miss | Split | Spurious | Accurate | Precision | Recall | F-Measure |
| Nessys | 0,68 | 0,97 | 91,55 | 0,14 | 5,76 % | 0,00 % | 3,66 % | 1,29 % | 89,30 % | 95,05 % | 94,24 % | 94,65 % |
| Ilastik | 0,53 | 0,96 | 92,21 | 0,33 | 27,30 % | 4,26 % | 23,45 % | 3,13 % | 41,86 % | 71,75 % | 68,85 % | 70,27 % |
| MINS | 0,41 | 0,95 | 92,11 | 0,52 | 28,94 % | 2,35 % | 19,75 % | 3,57 % | 45,39 % | 75,06 % | 69,95 % | 72,41 % |
| FarSight | 0,58 | 0,96 | 92,95 | 0,14 | 39,25 % | 0,00 % | 8,71 % | 19,58 % | 32,47 % | 70,42 % | 62,14 % | 66,02 % |

| Blastocysts (5 images - 699 Cells) |  |  |  |  |  |  |  |  |  |  |  |  |
| --- | --- | --- | --- | --- | --- | --- | --- | --- | --- | --- | --- | --- |
| Method | Jl | RI | Hausdorff | NSD | Merge | Miss | Split | Spurious | Accurate | Precision | Recall | F-Measure |
| Nessys | 0,69 | 0,96 | 80,37 | 0,17 | 0,17 % | 4,67 % | 2,57 % | 0,00 % | 92,59 % | 97,43 % | 95,15 % | 96,28 % |
| Ilastik | 0,73 | 0,97 | 80,80 | 0,16 | 1,17 % | 1,34 % | 4,75 % | 0,65 % | 92,10 % | 94,60 % | 97,48 % | 96,02 % |
| MINS | 0,71 | 0,96 | 81,34 | 0,21 | 3,51 % | 0,17 % | 4,68 % | 1,78 % | 89,87 % | 93,40 % | 96,33 % | 94,84 % |
| FarSight | 0,70 | 0,95 | 78,05 | 0,03 | 15,66 % | 0,00 % | 3,63 % | 1,83 % | 78,89 % | 94,55 % | 84,65 % | 89,32 % |

| E7.5 (2 Zones 924 Cells) |  |  |  |  |  |  |  |  |  |  |  |  |
| --- | --- | --- | --- | --- | --- | --- | --- | --- | --- | --- | --- | --- |
| Method | Jl | RI | Hausdorff | NSD | Merge | Miss | Split | Spurious | Accurate | Precision | Recall | F-Measure |
| Nessys | 0,59 | 0,87 | 283,01 | 0,17 | 6,27 % | 2,34 % | 4,16 % | 0,35 % | 86,88 % | 95,49 % | 91,44 % | 93,42 % |
| Ilastik | 0,70 | 0,90 | 272,92 | 0,09 | 6,34 % | 0,00 % | 6,86 % | 0,66 % | 86,14 % | 92,48 % | 93,64 % | 93,05 % |
| MINS | 0,54 | 0,87 | 283,50 | 0,47 | 14,71 % | 1,67 % | 22,69 % | 2,87 % | 58,07 % | 74,44 % | 84,10 % | 78,98 % |
| FarSight | 0,60 | 0,83 | 123,24 | 0,01 | 18,57 % | 0,00 % | 28,81 % | 2,71 % | 49,91 % | 68,48 % | 82,67 % | 74,91 % |

| E8.75 (3 zones - 4593 Cells) |  |  |  |  |  |  |  |  |  |  |  |  |
| --- | --- | --- | --- | --- | --- | --- | --- | --- | --- | --- | --- | --- |
| Method | Jl | RI | Hausdorff | NSD | Merge | Miss | Split | Spurious | Accurate | Precision | Recall | F-Measure |
| Nessys | 0,64 | 0,85 | 991,02 | 0,26 | 3,11 % | 3,03 % | 2,05 % | 0,81 % | 91,00 % | 97,14 % | 93,81 % | 95,45 % |
| Ilastik | 0,65 | 0,86 | 988,70 | 0,34 | 8,15 % | 2,92 % | 6,28 % | 1,28 % | 81,38 % | 92,44 % | 88,84 % | 90,61 % |
| MINS | 0,56 | 0,82 | 1004,26 | 0,46 | 13,10 % | 0,55 % | 16,29 % | 2,89 % | 67,17 % | 80,82 % | 86,55 % | 83,59 % |
| FarSight | 0,63 | 0,84 | 1033,03 | 0,18 | 21,30 % | 0,30 % | 7,11 % | 2,51 % | 68,79 % | 90,39 % | 78,43 % | 83,98 % |

**Table S3:** Description of the measures provided by the segmentation benchmarking tool.

| ClusteringComparator | Name of the java class generating the measures listed below |
| --- | --- |
| Jaccard Index | Similarity measure (higher values are better) |
| Rand Index | Similarity measure (higher values are better) |
| Hausdorff Mean | maximum of the set of minimal distances between two compared shapes (lower values are better) |
| Hausdorff Std | Standard deviation of the Hausdorff measure distribution |
| NSD Mean | Normalised Sum of distances : average distance of mis-labelled pixels by the automated method from the border of the reference shape in GT (lower values are better) |
| NSD Std | Standard deviation of the NSD measure distribution |
| SegmentationAccuracy | Name of the java class generating the measures listed below |
| Merge count | Number of merged events |
| Merge size Mean | Average number of shapes per merge event |
| Merge size Std | Stand deviation of the distribution of the number of shapes per merge events |
| Miss | Number of miss events |
| Ref Number of Border Cells | Number of shapes touching the image border in the GT image |
| Ref Total Number of Cells | Total number of shapes in the GT image |
| Split count | Number of split events |
| Split size Mean | Average number of shapes per split event |
| Split size Std | Standard deviation of the distribution of the number of shapes per split events |
| Spurious | Number of spurious detection occurrences |
| Stretch | Number of detected shapes in the tested image which overlap with more than one GT shape (at least 25 % of detected shape volume). This can encompass merged or split events |
| Test Number of Border Cells | Number of shapes touching the image border in the tested image |
| Tested Total Number of Cells | Total number of shapes in the tested image |

**Table S4:**
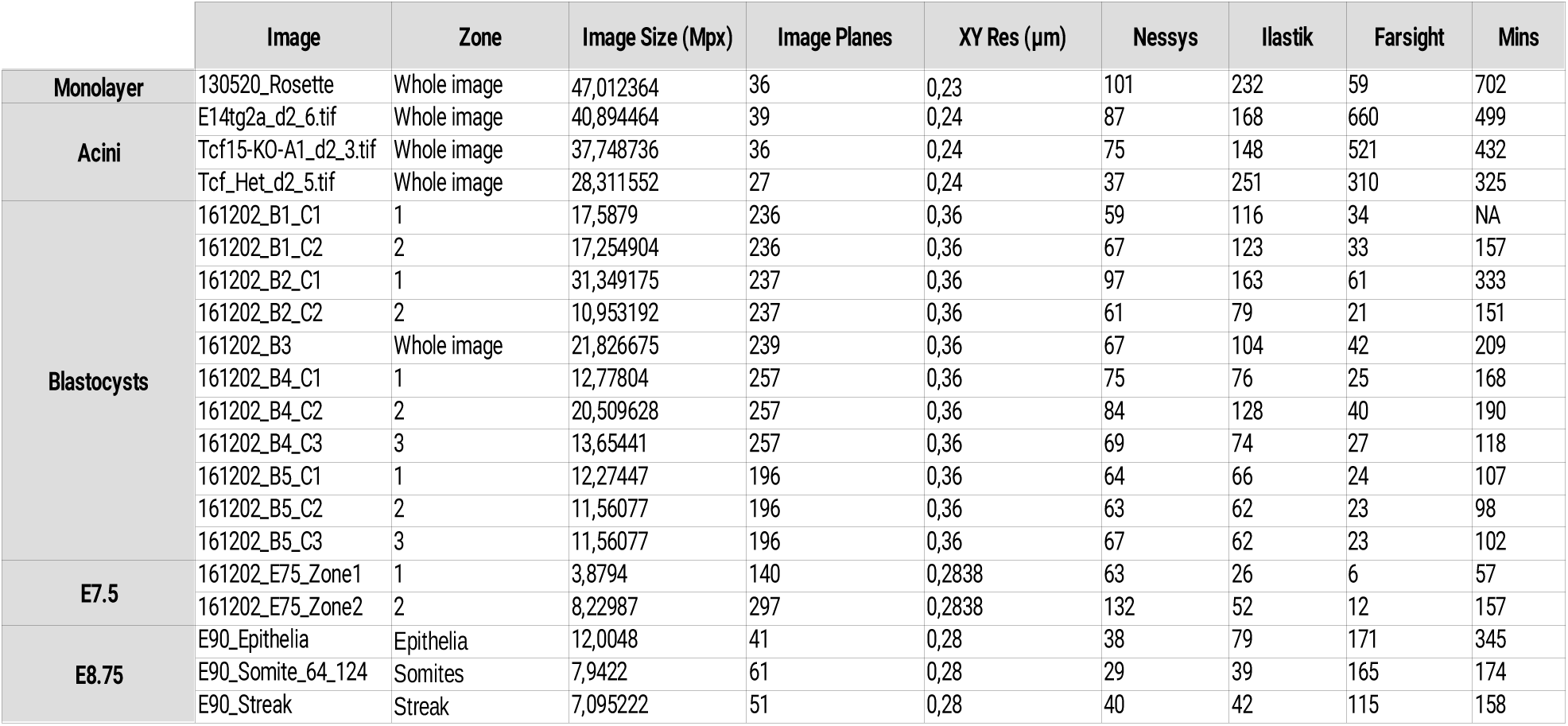
Processing Time for each image and method.

**Table S5:** Nessys processing time details with increasing image plane size or plane number.

|  | Image | Image Size |  |  | Image Properties |  |  | Processing time (Sec.) |  |  |  |  |  |
| --- | --- | --- | --- | --- | --- | --- | --- | --- | --- | --- | --- | --- | --- |
|  |  | X | Y | Z | Mpx | Cell Number | Cells/Mpx | Steer | RBS | Linkage | Total | Mvx/Sec | Cells/Sec |
| Increasing Plane Size | A1 | 472 | 359 | 145 | 24,56996 | 4511 | 183,6 | 3 | 105 | 21 | 129 | 0,190 | 35,0 |
|  | A2 | 665 | 505 | 145 | 48,694625 | 7956 | 163,4 | 6 | 172 | 34 | 212 | 0,230 | 37,5 |
|  | A3 | 940 | 715 | 145 | 97,4545 | 15280 | 156,8 | 12 | 276 | 63 | 351 | 0,278 | 43,5 |
|  | A4 | 1330 | 1011 | 145 | 194,97135 | 23542 | 120,7 | 24 | 428 | 104 | 556 | 0,351 | 42,3 |
|  | A5 | 1881 | 1430 | 145 | 390,02535 | 35443 | 90,9 | 48 | 649 | 165 | 862 | 0,452 | 41,1 |
|  | A6 | 2660 | 2022 | 145 | 779,8854 | 61540 | 78,9 | 102 | 1080 | 327 | 1509 | 0,517 | 40,8 |
|  | A7 | 3763 | 2860 | 145 | 1560,5161 | 81284 | 52,1 | 199 | 1550 | 502 | 2251 | 0,693 | 36,1 |
| Increasing Plane Number | B1 | 1024 | 1024 | 24 | 25,165824 | 4032 | 160,2 | 6 | 81 | 21 | 108 | 0,233 | 37,3 |
|  | B2 | 1024 | 1024 | 48 | 50,331648 | 7892 | 156,8 | 10 | 158 | 42 | 210 | 0,240 | 37,6 |
|  | B3 | 1024 | 1024 | 96 | 100,6633 | 15485 | 153,8 | 21 | 302 | 72 | 395 | 0,255 | 39,2 |
|  | B4 | 1024 | 1024 | 192 | 201,32659 | 30676 | 152,4 | 41 | 602 | 144 | 787 | 0,256 | 39,0 |
|  | B5 | 1024 | 1024 | 384 | 402,65318 | 61108 | 151,8 | 81 | 1270 | 281 | 1632 | 0,247 | 37,4 |
|  | B6 | 1024 | 1024 | 580 | 608,17408 | 92250 | 151,7 | 113 | 1827 | 556 | 2496 | 0,244 | 37,0 |

**Table S6:**
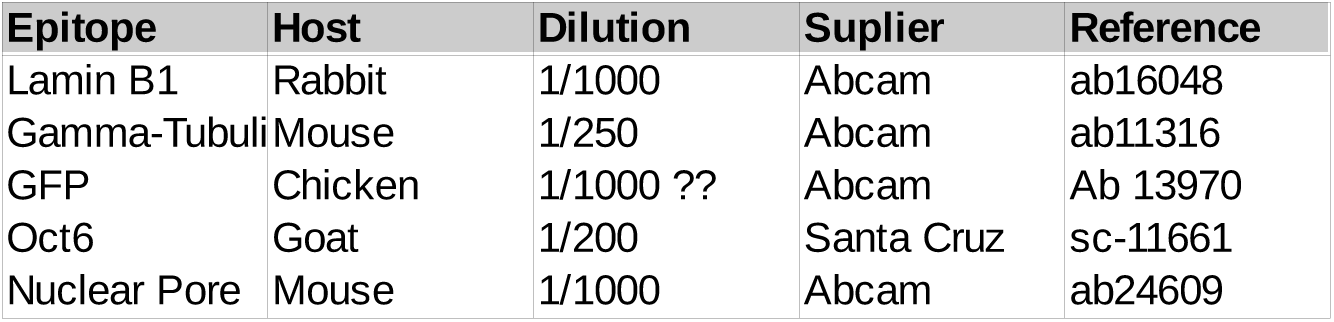
Primary antibodies used in this study.

| Epitope | Host | Dilution | Supplier | Reference |
| --- | --- | --- | --- | --- |
| Lamin B1 | Rabbit | 1/1000 | Abcam | ab16048 |
| Gamma-Tubulin | Mouse | 1/250 | Abcam | ab11316 |
| GFP | Chicken | 1/1000 ?? | Abcam | Ab 13970 |
| Oct6 | Goat | 1/200 | Santa Cruz | sc-11661 |
| Nuclear Pore | Mouse | 1/1000 | Abcam | ab24609 |

**Movie S1 Examples of Nessys Segmentation outputs** Plane by plane animation of the Nessys segmentation outputs shown in Fig. 1B and C. The LaminB1 signal is overlaid in cyan.

**Movie S2 Illustration of the Ridge Following Procedure** This movie is a slow motion of the ridge following procedure. The movie starts by a display of the final tree as described in the legend of figure 2. The green portion of the tree represents the pair of leaves selected as the most probable valid shape according to the ranking performed by the classifier. Then a sequence shows how the tree was grown. The green dot represents the location of the ridge following procedure at a given time and the blue lines highlight the ridges that have already been identified. Finally the leaves, the root and the ‘winning’ ridge are highlighted before the segmented area is drawn.

**Movie S3 Time lapse of double colour reporter cells during neural differentiation**. Movie of the time lapse experiment shown in Fig. 7. Green: NE-mKate2 signal, Red: Sox1-GFP. 1 sec = 38 min in real time.

